# β2 adrenergic receptors orchestrate neutrophil demargination and recruitment to the ischemic heart following myocardial infarction

**DOI:** 10.1101/2025.10.20.683429

**Authors:** Albert Dahdah, Ki Ho Park, Krishna P Maremanda, Mathankumar Marimuthu, Robert M. Jaggers, Dipanjan Chattopadhyay, Wang Qiang, Nitin Nitin, Stavros Stavrakis, Tarun W. Dasari, Viorel I. Suicã, Raluca M. Boteanu, Elena Uyy, Felicia Antohe, Douglas G. Tilley, Andrew J. Murphy, Prabhakara R. Nagareddy

**Author notes:** **Corresponding author,** Prof. Prabha Nagareddy, PhD, FAHA, 1122 NE, 13th Street, Suite 333B, Oklahoma City OK 73117, USA.

## Abstract

Neutrophils play a crucial role in instigating inflammation as well as its resolution post-myocardial infarction (MI). Although granulopoiesis in the bone marrow (BM) is the major source of cardiac neutrophils post-MI, infiltration of neutrophils to the heart occurs much quicker than peak granulopoiesis. These observations suggest that sources other than granulopoiesis may supply neutrophils to the heart during the early hours post-MI. Using a combination of flow cytometry, BM ablation of hematopoietic stem cells, confocal microscopy and multiple proteomics analysis, we found that the first wave of neutrophils recruited to the ischemic heart is exclusively sourced from vasculature and not from granulopoiesis in the BM/ spleen. The MI-evoked neutrophilia during the early hours bore all hallmarks of demargination induced by classical demarginating agents such as dexamethasone/ norepinephrine (NE). Various pharmacological and genetic strategies aimed at suppressing NE synthesis or disruption of β-AR signaling reduced both neutrophil demargination as well as recruitment to the heart. Interestingly, however, despite a marked reduction in cardiac neutrophil burden only short-term inhibition of β-ARs improved cardiac remodeling and function. Our findings support a pharmacological strategy to contain the initial onslaught of neutrophils on the ischemic heart using β2-AR blockers to regulate the otherwise runaway inflammatory response.

## Introduction

Acute myocardial infarction (AMI) induces a rapid and robust inflammatory response characterized by infiltration of different immune cell types to the infarcted heart. The recruitment of immune cells is a dynamic process and consists of sequential infiltration of neutrophils, pro- and anti-inflammatory monocytes, and lymphocytes to the injured myocardium^1-6^. While the excessive inflammatory response after MI is responsible for cardiomyocyte death, infarction expansion and adverse remodeling, it also has an essential role in orchestrating the reparative response for optimal infarct healing^7^. The inflammatory response following MI can be mapped into three distinct but overlapping phases^8-11^. The first phase, also known as the acute inflammatory phase, occurs at the onset of MI and is characterized by a significant increase in innate immune cell (neutrophils as well as monocytes/ macrophages) infiltration to the myocardium mainly to clear debris and dead cells. In the second phase, reparative monocytes, macrophages, dendritic cells, and lymphocytes are mobilized to resolve the initial inflammatory response and support the formation of vascularized granulation tissue (from 3-4 days to 2 weeks after infarction). The third phase takes place from 2 weeks onward where the vascularized granulation leads to a stable myocardial scar that is almost free of immune cells. It is therefore important that a timely and well-balanced inflammatory response is established to successfully dampen the inflammatory response and initiate infarct healing in patients surviving MI. However, there are no clear strategies in clinical practice to regulate the inflammatory response optimal enough to favorably modulate the remodeling processes.

Studies from our lab and elsewhere have shown that infiltration of neutrophils to the infarcted myocardium have the ability to influence the nature of the ensuing inflammatory response. This is because neutrophils are the first responders to ischemic injury and have the ability to amplify the initial inflammatory response by deploying key alarmins such as S100A8 and S100A9. These alarmins interact with Toll-Like-Receptor (TLR4) on naïve neutrophils, prime the NLRP3 inflammasome and facilitate their migration to the bone marrow (BM). The reverse-migrated neutrophils are preferentially retained within the BM sinusoids by certain adhesion receptors, a process that is essential for the activation of Gasdermin D-dependent membrane pore formation and secretion of IL-1β^12,13^. All these events culminate in amplified granulopoiesis and ensure a steady supply of neutrophils to facilitate the clearance of debris and dead cells and, to initiate wound healing. Interestingly, the process of granulopoieses in the BM does not reach the climax until 24 hours post-MI, yet neutrophils begin to appear in the heart much faster than their production rate in the BM. These findings suggest that sources other than granulopoiesis may help replenish neutrophils for immediate deployment to the ischemic heart. However, the exact source of these neutrophils is unclear.

Under steady-state conditions, mature neutrophils are present in major reservoirs across the body (spleen, liver, BM, lungs, and vasculature) and are recruited following changes in homeostasis^14^. The unique feature of neutrophils in the vasculature is their distribution into two distinct groups, the marginated and the demarginated. Firmly adherent to the vasculature walls, the marginated pool of neutrophils represents ∼ 51% of neutrophils in circulation and are believed to serve as early responders. The demargination process in neutrophils is attributed to loss of adhesion molecules, and rearrangement of actin cytoskeleton within the cells^15,16^. Under certain stress and inflammatory conditions, adhesion of neutrophils is altered, and the marginated cells detach and mobilize into the bloodstream, expanding the percentage of demarginated cells in the blood^17^. We reasoned that the marginated pool of neutrophils may be mobilized to meet the sudden and increased demand of the ischemic heart. Thus, deciphering the signaling pathways that stimulate the demargination of neutrophils and their recruitment to the ischemic heart may provide a viable approach to regulate neutrophil trafficking to the heart post-MI.

## Methods

Detailed methods including statistics are provided in the online-only Data Supplement. All animal studies were approved by the IACUCs in accordance with the NIH Guide for the Care and Use of Laboratory Animals. Detailed analytical methods and reagents will be made available to other investigators on a reasonable request. Key resources and reagents used in the study are listed in Table-S1.

### Statistics

Normal distribution was evaluated using the Shapiro-Wilk test. For statistical comparisons of 2 groups, an unpaired t test (for normally distributed variables), or a Mann-Whitney test (for non-normally distributed variables) was applied. For comparing 3 or more groups with normal distribution data, a one-way or two-way analysis of variance (ANOVA) followed by either a Holm-Sidak’s or Tukey’s multiple comparison tests was applied. For non-normal data, a Kruskal-Wallis test followed by a Dunn’s multiple comparisons test was applied. Comparisons of the survival endpoint between treatment groups were performed with the log-rank test. When data were collected over time on the same set of animals such as echocardiography parameters, they were analyzed using Two-Way ANOVA (mixed-effects model) followed by Bonferroni’s multiple comparison test. A P value ≤ 0.05 was used as a cut off for statistical significance throughout the analyses. All analyses were done using GraphPad Prism version 8.

## Results

### Neutrophil recruitment to the ischemic heart precedes the peak of bone marrow (BM) granulopoiesis

We previously have shown that large-scale infiltration of neutrophils to the ischemic heart post-MI^12^ is facilitated by a robust activation of HSPCs in the BM (∼18-24 hours). Furthermore, the first wave of recruited neutrophils releases alarmins (e.g., S100A8 and S100A9) via NETosis, a crucial step in sustaining granulopoiesis in the BM^13^. However, the pathways that regulate granulopoiesis in the BM are activated between 12- and 24-hours post-MI and thus, the mechanisms that trigger the recruitment of the first wave of neutrophils are unknown. We hypothesized that the circulating neutrophils including the marginated pool are sourced to meet the immediate demand of the ischemic heart. To test this hypothesis, we first performed a temporal study to examine the kinetics of neutrophil recruitment to the heart and their production from sites in the BM and spleen. We employed a mouse model of permanent ligation of the left anterior descending (LAD) artery as described previously^12,13^.

Following MI, an increase in neutrophil recruitment to the heart occurred swiftly and reached a submaximal threshold as around 6 hours (Figure. 1A). These heart-infiltrating neutrophils were likely sourced from circulation since blood also showed a significant increase in neutrophil number at 6 hours after MI (Figure 1B). The neutrophils are matured and not newly produced because the EdU (a nucleoside analog of thymidine incorporated into DNA during active DNA synthesis) label did not increase significantly in neutrophils in both the heart (Figure 1C) and blood (Figure 1D) up until 24 hours post-MI. We also confirmed that the increase in neutrophil number in the heart and blood during the first 6 hours was not due to mobilization of matured neutrophils from the BM (Figure 1E) since the neutrophil stocks in BM remain undepleted (45% vs 37%). Interestingly, the number of neutrophils in the spleen increased (Figure 1F). However, as expected, the EdU label did not increase in either BM (Figure 1G) or splenic neutrophils (Figure 1H) suggesting a secondary role for granulopoiesis during the initial hours after MI. This was also reflected in BM progenitor cells (i.e. CMPs, GMPs) that did not show any sign of enhanced granulopoiesis up until 24 hours post-MI (Figure 1,I to K). This led us to conclude that the initial wave of neutrophils recruited to the heart (< 6 hours post-MI) was not sourced from the BM/ splenic reservoirs or granulopoiesis (Summarized in Figure 1L).

**Figure 1.**
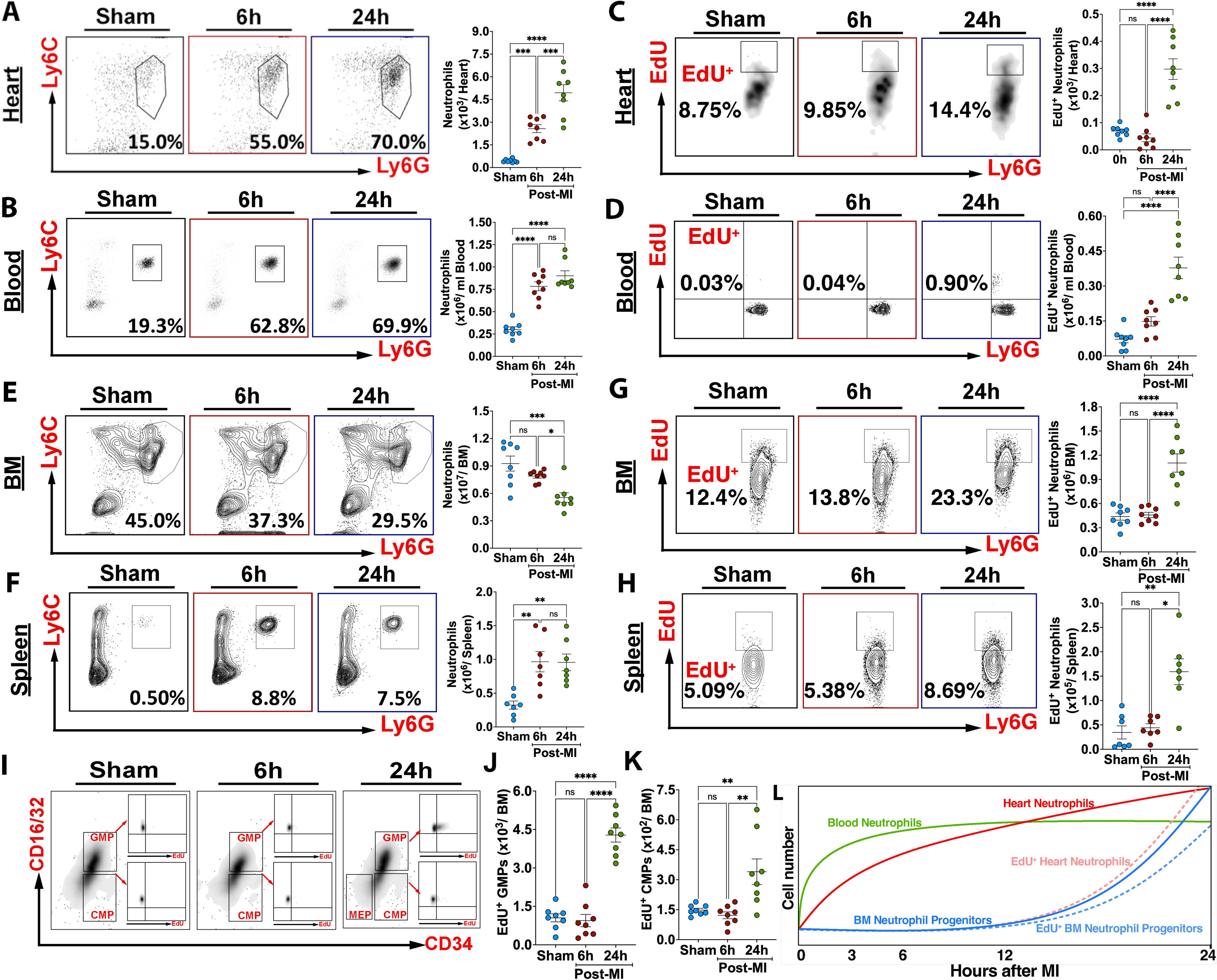
Bone marrow (BM) granulopoiesis is not the primary source of cardiac neutrophils during the early hours following myocardial infarction (MI). C57BL/6J WT mice were subjected to MI by permanent ligation of the left anterior descending coronary (LAD) artery. Blood, heart, bone marrow (BM) and spleen tissues were collected at various time points as indicated in the individual figures to assess the abundance of neutrophils and their progenitors (BM) by flow cytometry. Non-MI groups served as sham controls. Representative flow cytometric plots (left panel) and quantification (right panel) of matured neutrophils in the heart (A) and blood (B). Representative flow cytometric plots (left panel) and quantification (right panel) of newly produced (EdU^+^) neutrophils in the heart (C) and blood (D). Representative flow cytometric plots (left panel) and quantification (right panel) of matured neutrophils in the BM (E) and spleen (F). Representative flow cytometric plots (left panel) and quantification (right panel) of newly produced (EdU^+^) neutrophils in the BM (G) and spleen (H). I, Flow cytometric gating (left panel) and quantification (right panel) of proliferation rates of granulocyte macrophage progenitors, GMPs (J) and common myeloid progenitors, CMPs (K) in the BM. L, A schematic illustration (not to scale) summarizing neutrophil kinetics and granulopoiesis over a period of 24 hours post-MI. In response to MI, matured neutrophil number increases rapidly in blood and heart while the newly produced neutrophils (EdU^+^) are observed in the blood and heart only during peak granulopoiesis in the BM (∼18-24 hours). All data are means ± SEM. Statistical tests for A to G and K (1-way ANOVA and Tukey’s post-hoc test, *P<0.05, **P<0.001, ***P<0.0005, ****P<0.0001) while H (Kruskal Wallis+ Dunn’s, *P<0.05, **P<0.005, n.s, not significant). All cells were gated after removing debris, dead and clustered cells. Neutrophils were gated from CD45^+^ cells. CMPs and GMPs were gated from myeloid progenitor cells (Lin^-^, ckit^+^, Sca1^-^). Newly generated neutrophils and granulopoiesis in the BM was assessed based on the incorporation of EdU label. The number in the parentheses in A to H indicates the neutrophil population as a percentage of CD45^+^ cells while in C, D, G,H, J and K the number indicates the percentage of EdU^+^ neutrophil populations.

### Neutrophil demargination from the vasculature is the primary source of neutrophils in the ischemic heart during the early hours following MI

Since we found no major role for either BM or spleen in supplying neutrophils to the ischemic heart immediately after-MI, we hypothesized that neutrophils within the vasculature are the predominant source. To verify this notion, we performed a series of experiments utilizing the total body irradiation (TBI) strategy along with the administration of classical demarginating agents such as dexamethasone (DEX) and nor-epinephrine (NE). We reasoned that TBI of mice would lead to complete ablation of proliferating HSPCs and, thus granulopoiesis in the BM/ spleen would be severely suppressed. If our hypothesis is true, administration of DEX or induction of MI post-TBI would still increase the number of demarginated neutrophils in the absence of routine granulopoiesis.

First, to confirm that DEX and NE can be used as experimental agents, we performed a temporal study. We injected healthy WT mice with a single dose of either NE or DEX and quantified the number of neutrophils demarginated over a period of 24 hours. Using flow cytometry, we quantified the neutrophil numbers in blood and found a rapid increase starting at 2 hours and reaching the peak at 4 hours following NE administration, while the neutrophil population in DEX-treated mice reached a maximum at around 8 hours (Supplemental Figure 1A). These data suggest that both NE and DEX induce demargination but may follow different trajectories perhaps due to different mechanisms of demargination. Next, to confirm that TBI does prevent granulopoiesis without adversely impacting matured blood neutrophils, healthy WT mice were lethally irradiated (TBI) following which the number of neutrophils in the blood and their progenitor cells in the BM and spleen were analyzed over a period of 24 hours. TBI of mice led to a significant decrease in the number of progenitor cells in the BM (Supplemental Figure 1B) and spleen (Supplemental Figure 1C) by 6 hours while the number of matured neutrophils in the blood (Supplemental Figure 1D), BM (Supplemental Figure 1E) and spleen (Supplemental Figure 1F) began to decline after 6 hours and remained constantly lower thereafter. The decline in the number of blood neutrophils can be attributed to the lack of normal hematopoiesis and replenishment due to radiation damage caused to HSPCs. After initial characterization of neutrophil response separately to TBI and demarginating agents, we selected a time point (∼18 hours) wherein the blood neutrophil count is slightly decreased, and the BM is unable to mount a granulopoietic response (Summarized in Figure 2A). We choose this time point as a window of opportunity to study the impact of DEX or MI on neutrophil response. As expected, administration of DEX to irradiated mice (Study outline, Figure 2B) significantly increased the number of neutrophils in the blood (Figure 2C) despite a dramatic depletion of progenitor cells in the BM and spleen (Supplemental Figure 1, B and C). Similarly, induction of MI in TBI mice (Study outline, Figure 2D) also increased the number of neutrophils in the infarcted heart (Figure 2E) given that a significantly elevated number of demarginated neutrophils was available in the blood (Figure 2F) for rapid recruitment to the heart. This increase in the number of circulating neutrophils indicated that a population of neutrophils is actively present and remains marginated in the vasculature until certain stress signals induce their demargination.

**Figure 2.**
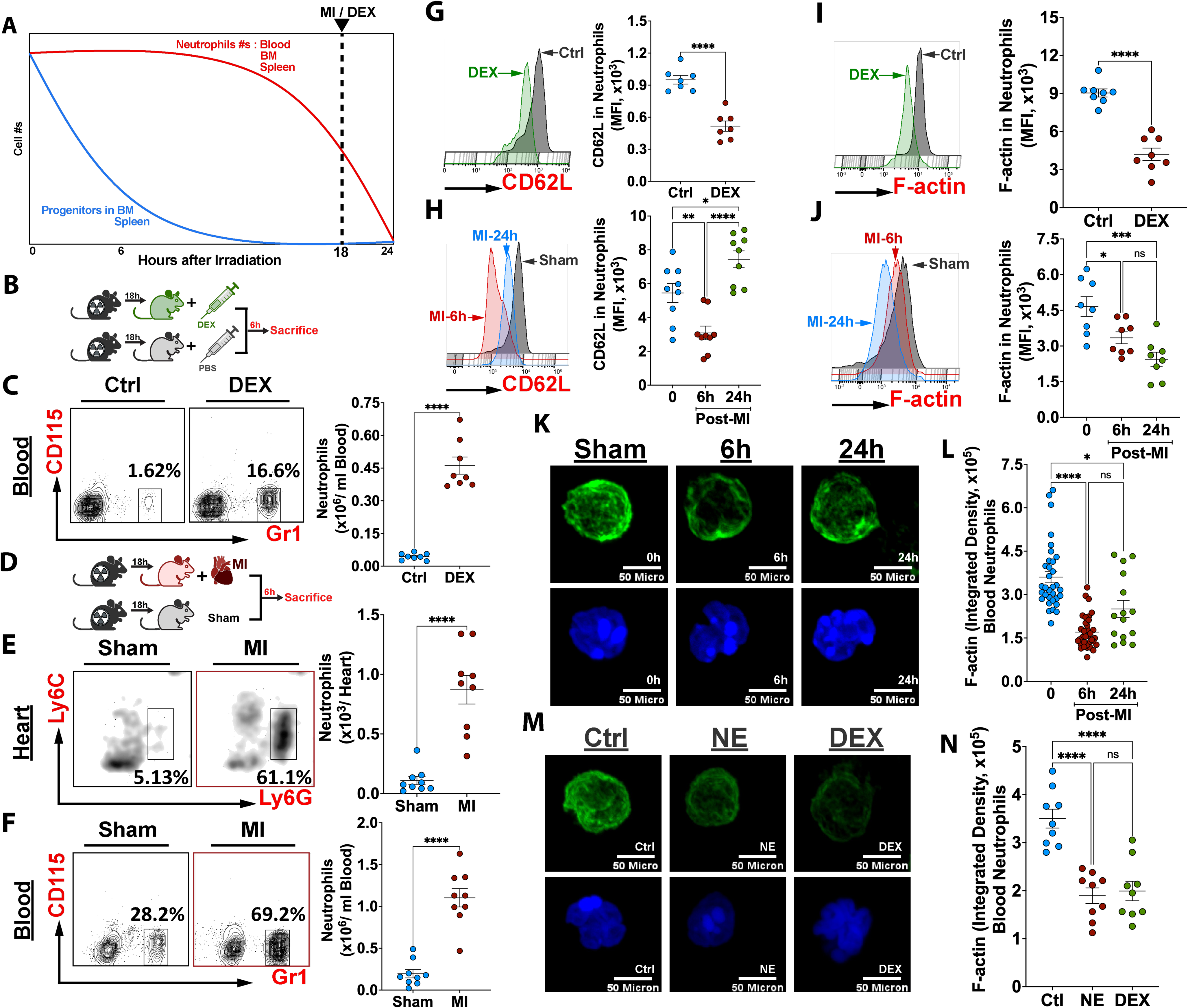
Vascular demargination is the predominant source of cardiac neutrophils during the early hours following MI. **A,** Schematic illustration (not to scale) depicting chronological depletion of neutrophils and their progenitors from various tissue compartments over a period of 24 hours following total body irradiation (TBI). By 24 hours, all animals become extremely neutropenic as a result of total ablation of HSPCs in the BM and spleen. The time point of 18 hours post-TBI was chosen to intervene with either MI or demarginating agents (DEX) to identify the source of cardiac neutrophils post-MI. **B**, A schematic figure depicting the experimental strategy to study the effect of DEX (administered at 18 hours post-TBI) on neutrophil demargination. **C**, Representative flow cytometric plots (left panel) and quantification (right panel) of neutrophils in the blood at 6 hours after DEX administration. **D**, A schematic figure depicting the experimental strategy to study the impact of MI (induced at 18 hours post-TBI) on neutrophil demargination. **E**, Representative flow cytometric plots (left panel) and quantification (right panel) of neutrophils in the heart at 6 hours post-MI. **F**, Representative flow cytometric plots (left panel) and quantification (right panel) of neutrophils in the blood at 6 hours post-MI. **G**, Representative histograms (left panel) depicting cell surface expression of CD62L and quantification (right panel) of mean fluorescence intensity (MFI) on neutrophils in the blood of control and DEX treated mice. **H,** Representative histograms (left panel) depicting cell surface expression of CD62L and quantification (right panel) of MFI in neutrophils in the blood of sham and MI mice. **I**, Representative histograms (left panel) depicting F-actin stain intensity and quantification of MFI (right panel) on neutrophils in the blood of control and DEX treated mice. **J**, Representative histograms (left panel) depicting F-actin stain intensity and quantification (right panel) of MFI in neutrophils in the blood of sham and MI mice. **K**, Representative confocal images of demarginated neutrophils stained with F-actin and DAPI at 0, 6- and 24-hours post-MI and (**L**) quantification of F-actin intensity. **M**, Representative confocal images of demarginated neutrophils stained with F-actin and DAPI at 6 hours following NE or DEX administration and (**N**) quantification of F-actin intensity. All data are means ± SEM, Statistical tests for **C, E, F, G and I** (Unpaired t test, ****P<0.0001), **H, J** and **N** (1-way ANOVA and Tukey’s post-hoc test, *P<0.05, **P<0.001, ***P<0.0005, ****P<0.0001, n.s, not significant), **L** (Kruskal Wallis+ Dunn’s, ****P<0.0001, n.s not significant). Neutrophil numbers in the blood, heart, and staining intensity (i.e. MFI) of CD62L and F-actin were quantified using flow cytometry. MI, Myocardial infarction; TBI, Total body irradiation; DEX, Dexamethasone; NE, Nor-epinephrine.

To confirm that demarginated neutrophils are the predominant source of heart infiltrating neutrophils during the early hours after-MI, we tested an alternate hypothesis wherein we first induced MI in two groups of healthy WT mice, followed by injection of DEX (Study outline, Supplemental Figure 1G). We reasoned that administration of DEX to MI mice would not further increase the number of neutrophils in the blood or heart. This is because a prior MI would have exhausted all the marginated neutrophils, and there are no more neutrophils left for DEX to demarginate them. As expected, injection of DEX to MI mice did not increase the number of neutrophils in the blood (Supplemental Figure 1H) or the heart (Supplemental Figure 1I) confirming that there were no more marginated neutrophils left in the vasculature.

Following MI, matured neutrophils can be sourced from the BM or spleen by emptying, a process termed as forced mobilization. To assess their contribution to the demarginated pool and their potential recruitment to the heart, we induced MI in a separate group of mice and immediately injected granulocyte colony-stimulating factor (G-CSF) (Study outline, Supplemental Figure 1J). As expected, the administration of G-CSF induced a rapid efflux of neutrophils (perhaps premature) from the BM with a proportional decrease in BM, a response that is typical to G-CSF administration (Supplemental Figure 1K). These data suggest that MI did not mobilize/ empty the BM of neutrophils during the first 6 hours. Interestingly, the increase in the number of circulating neutrophils (Supplemental Figure 1L) in response to G-CSF did not translate into greater cardiac neutrophil burden (Supplemental Figure 1M). These data not only highlight the importance of the source of neutrophils (i.e., demargination vs BM emptying) but also confirm that matured neutrophils from vasculature are preferred over immature cells from the BM for recruitment to the infarct.

### Following MI, neutrophils display phenotypic hallmarks characteristic of demarginated neutrophils

Constitutively expressed adhesion molecules such as P- and L-selectin (CD62L) are important for neutrophils to remain anchored to the vascular endothelium. During neutrophil migration (i.e., demarginated) and trafficking into injured tissues, CD62L is rapidly shed. It is long known that anti-inflammatory glucocorticoids (e.g., DEX) and catecholamines (e.g., NE) induce neutrophil demargination and, thus neutrophilia, by suppressing CD62L gene expression and/ or shedding of surface CD62L by Adam17, a sheddase^16,18^. Recent studies have shown that, in addition to the shedding of CD62L, a purely mechanical phenomenon caused by cell softening also plays a major role in neutrophil demargination^19^. Exposure of neutrophils to DEX or NE reorganizes cellular cortical actin (i.e., reduction in localized and polymerized actin), significantly decreasing cell stiffness and thus, allowing them to demarginate. To confirm if blood neutrophils in MI mice exhibit demargination signatures similar to that induced by DEX or NE treatment, we measured the cell surface expression of CD62L and F-actin intensity by flow cytometry and confocal microscopy respectively. Interestingly, neutrophils in both MI and DEX/ NE treated mice showed strikingly similar reduced expression of CD62L (Figure 2, G and H) and F-actin intensity (Figure 2, I-N). The untreated neutrophils reveal a much brighter, continuous cortical polymerized actin ring and high amounts of localized actin, while the MI (6 hours) and NE-treated neutrophils show a much dimmer but still continuous polymerized actin ring. DEX-treated neutrophils were highly heterogeneous with discontinuous cortical polymerized actin (Figure 2, M and N). It is important to note that the surface expression of CD62L was decreased in MI neutrophils at around 6 hours (typical demargination) but began to increase thereafter (i.e., 24 hours) (Figure 2H). These observations suggest that the neutrophils in blood at 24 hours post-MI are newly produced via granulopoiesis in the BM. Together, these findings support our hypothesis that MI promotes neutrophil demargination via mechanisms that involve CD62L shedding and cell softening by stimulating the reorganization of the cortical actin cytoskeleton. We then turned to exploring how neutrophils demarginate and are recruited to the heart.

### Proteomic analysis of demarginated neutrophils after MI or NE administration reveals activation of pathways involved in cellular movement and cytoskeletal reorganization

Neutrophils undergo transcriptional and translational changes while in the circulation and during their recruitment to injured tissues^13^. Having confirmed that the demarginated neutrophils contribute to increased neutrophil burden in the heart post-MI, we next focused on identifying the signaling pathways that promote demargination of neutrophils. To achieve this, we induced MI in healthy WT mice and sorted blood neutrophils at 6 hours post MI and subjected the samples to proteomics (Study outline, Figure 3A) using a high-resolution LC-MS/MS (label-free spectral counting approach). Purified neutrophils harvested from the blood at 6 hours after sham surgery served as controls (Figure 3B).

**Figure 3.**
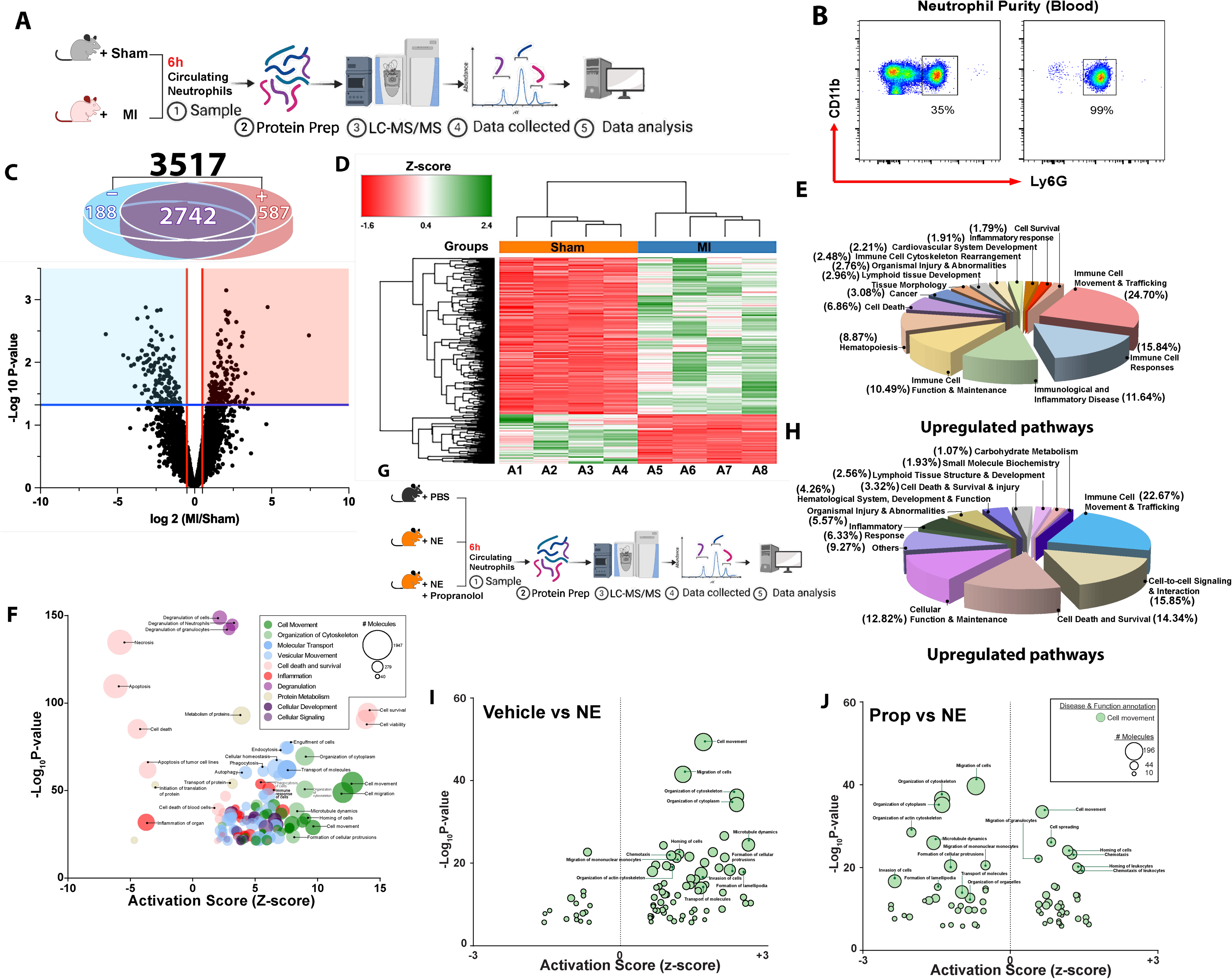
Pathways related to cellular movement and migration are enriched in demarginated neutrophils following MI or norepinephrine (NE) administration. **A**, A schematic figure depicting the experimental outline and workflow of neutrophil proteomics using a high-resolution LC-MS/MS (label-free spectral counting approach). MI was induced in healthy WT C57BL/6J mice and at 6 hours post-MI, neutrophils from peripheral blood were sorted using magnetic beads and purity tested by flow cytometry (**B**) following which they were subjected to whole cell proteomics analysis. Mice receiving sham surgery served as controls. **C**, A Venn diagram (top panel) showing the expression of proteins in demarginated neutrophils differentially impacted by MI compared to sham controls. **+** and **-** signs indicate the number of proteins upregulated and downregulated respectively by MI. The Volcano plot (bottom panel) revealing 775 differentially abundant proteins (DAPs) with a 1.5-fold change in abundance and an adjusted P value < 0.05, when MI group was compared to the Sham group. In total, there were 3517 identified high confidence proteins, identified based on a Sequest score > 10, a protein FDR < 0.01 and a minimum of 2 unique peptides. The red box contains the up-regulated proteins (587 in number), the blue box includes the down-regulated proteins (188 in number), and the remaining points on the white background indicate the proteins that were not significantly altered. **D.** Multivariate statistical hierarchical clustering heat map analysis of the MI and Sham proteomes depicting the proteins upregulated (green) or downregulated (red) in demarginated neutrophils in response to MI. Each column represents sorted blood neutrophils from individual mouse within MI (n=4, A5-A8) or Sham (n=4, A1-A4) groups. Proteins were triaged based on a p-value < 0.05 independent from their fold change values. **E**, Pie chart representing the pathways most impacted by MI. The number in the parentheses constitute the percentage representation of each pathway activated in response to MI. **F**, Visual representation of disease and function analysis using IPA; comparison and correlation between MI stimulated and unstimulated (sham) neutrophils to identify directions of change in major functions. Shown are the pathways upregulated in neutrophils by MI compared to sham mice. The extent of increase in each major function was assessed by plotting the activation scores against the possibility of occurrence. **G**, A schematic figure depicting the experimental outline and workflow of neutrophil proteomics. NE was administered to mice in the presence or absence of propranolol injected 8 hours prior to injection of NE. An additional dose of propranolol was injected along with NE. Six hours after administration of NE, demarginated neutrophils from peripheral blood were sorted using magnetic beads and subjected to proteomics. **H**, Pie chart representing the pathways most impacted by NE. The number in the parentheses constitute the percentage representation of each pathway activated by NE. **I**, IPA downstream effect analysis; comparison and correlation between NE stimulated and unstimulated neutrophils to identify the directions of change in major functions. Shown are the pathways upregulated by NE compared to vehicle treated mice. The extent of increase in the major functions was assessed by plotting the activation scores of the functions against the possibility of occurrence. **J**, Comparison, and correlation between NE stimulated neutrophils in the presence/ absence of propranolol to identify the directions of change in major functions. Shown are the pathways suppressed by propranolol compared to NE treated mice. The extent of increase in the major functions was assessed by plotting the activation scores of the functions against the possibility of occurrence. The Wilcoxon rank-sum test was used to determine whether proteins were differentially expressed between vehicle vs NE vs Prop groups. All statistical tests were two-sided, and statistical significance was considered when p value <0.05. To account for multiple-testing, the p values were adjusted using the Benjamini-Hochberg FDR correction. MI, Myocardial infarction; NE, Nor-Epinephrine; Prop, Propranolol.

MI induced a dramatic shift in neutrophil proteome as demonstrated by PCA plot depicting a clear spatial discrimination of the proteomes between the two groups (Supplemental Figure 2A). A total of 3517 protein entries were identified (Figure 3C). Among them, 587 proteins were upregulated, and 188 proteins were downregulated (with a 1.5-fold difference) in response to MI compared to sham group. A heat map view of the impacted proteins highlights the distinctive quantitative features between the 2 groups as well as the homogeneity of the biological replicates within each group (Figure 3D). Many of the proteins in demarginated neutrophils are unique to MI with dormant proteins flaring up compared to resting neutrophils in sham mice. Hierarchical pathway cluster analysis of the differentially abundant proteins revealed enrichment of various cellular processes and pathways unique to MI group (Supplemental Figure 2, B-G). Among the pathways most impacted by MI (Figure 3E) are those that are associated with cellular movement and immune cell trafficking (24.7%). An equal or slightly lower number of upregulated proteins are involved in cellular functions such as immune cell responses (15.8%), immunological/ inflammatory disease, (11.6%), immune cell function, and maintenance (10.5%). Delving deeper into different categories, pathways were then attributed to activation scores based on the level of combined altered functions of its constituent proteins and their effect in modifying neutrophil functions in MI (Figure 3F). We found a large number of proteins related to enhanced immune cell movement, migration, and organization of the cytoskeleton. Interestingly, the initial stages of MI also promoted cell survival, immune functions (phagocytosis) as well as a general increase in the alert state of an inflammatory response, while shutting down most pathways that are involved in the resolution.

Signaling pathways activated in neutrophils during an event such as MI are governed by a large number of ligands. Among the major canonical pathways that were activated and involved in neutrophilic response during MI, we found important modifications of proteins involved in G-Protein Coupled Receptor (GPCR) signaling. Signaling through GPCR from different ligands was found to converge in downstream actin polymerization, and cytoskeleton reorganization especially through the calcium signaling pathways^20^. These include Rac, Pak, Rho, as well as proteins involved in polarization and depolarization such as ROCK, COFILIN (CFN), PROFILIN, and Lim Kinase (LIMK) protein^21^ (Supplemental Figure 2H). For instance, the downstream Gα12/13 signaling of GPCRs upregulated RhoGEF and increased ROCK1 and ROCK2 that in turn block the myosin light chain kinase phosphatase (MLCP) and inhibit the phosphorylation of MLC proteins resulting in increased activation of phosphorylated myosin complexes that favor cytoskeleton reorganization^22^. ROCK proteins also increase the phosphorylation of LIMK, inhibiting phosphorylated active forms of CFN proteins responsible for actin filament stabilization^23^. On the other hand, downstream signaling through the GTPase Binding Domain (GBD) and the FYVE, RhoGEF and PH domain-containing proteins (FGD1/3) increase the serine/ threonine-protein Kinase (PAK), and the Wiskott–Aldrich Syndrome protein (WASP) that play a role in a variety of different signaling pathways involved in cytoskeleton regulation, cell motility, actin polymerization and actin filament reorganization^24,25^. Additionally, FGD1/3 is also known to activate the plasma-membrane associated small GTPase, the cell division cycle 42 (CDC42) proteins^26,27^. In its active state, CDC42 binds to a variety of effector proteins to regulate cellular attachment and actin polymerization^28,29^. We also found a slight upregulation of the glucocorticoid receptor NR3C1, however, no effective activation was predicted. In summary, the major impact of MI on signaling in neutrophils was found to converge on those pathways that typically represent actin polymerization, cytoskeletal reorganization, and cellular movement, downstream of GPCRs.

### Activation of β-adrenergic receptors by NE induces cellular movement and cytoskeletal reorganization, mirroring the effects observed following MI

Sympathetic response driven by β-adrenergic receptors (β-AR) constitutes the major GPCR signaling pathway in neutrophils^30^. To test the hypothesis that catecholamine stress induced by MI is the prime initiator of demargination and migration of neutrophils, we performed another proteomics study. Here, we treated healthy WT mice acutely with NE in the presence or absence of propranolol, a non-selective β-adrenergic receptor blocker (Study outline, Figure 3G). Administration of NE resulted in the activation of an immune response that was very similar to that induced by MI. Although we did not formally quantify the overlap in differentially expressed proteins between MI and NE groups, both conditions activated immune-related pathways—particularly those involved in cell movement and trafficking to a comparable extent (MI: 22.6; NE: 24.7), suggesting a similar overall biological response at the pathway levels. These include pathways representing immune cell movement and trafficking, cellular function, maintenance, and inflammation (Figure 3H). Among the pathways examined, the highest upregulated clustering was associated with cellular movement, organization of cytoplasm and actin cytoskeleton (Figure 3I). Most importantly, the majority of pathways associated with cell migration/ movement activated by the administration of NE in demarginated neutrophils were blocked by propranolol suggesting that the interaction between NE and β-adrenergic receptors is the main trigger for neutrophil demargination during MI (Figure 3J).

To define the specificity of the interaction between NE and β-ARs, we used various pharmacological agents to either prevent the synthesis of NE or its binding to one of the β-ARs. Experimental strategies include treatment of WT mice with a tyrosine hydroxylase inhibitor (Alpha-Methyl-p-Tyrosine (AMPT), non-selective β-AR blocker (propranolol), a selective β1 blocker (metoprolol) or a selective β2 blocker (butoxamine). Inhibition of tyrosine hydroxylase with AMPT (Study outline, Figure 4A) significantly suppressed neutrophil demargination (Figure 4B) and thus their subsequent recruitment to the heart (Figure 4C). Among the β AR blockers, the administration of each of the above agents separately produced a differential response. For example, treatment of mice with propranolol (Study outline, Figure 4D) suppressed both demargination (Figure 4E) and neutrophil recruitment to the heart (Figure 4F). However, blockade of β1 ARs (Study outline, Figure 4G) showed no impact on either neutrophil demargination (Figure 4H) or recruitment to the heart (Figure 4I) while inhibition of NE binding to β2 ARs (Study outline, Figure 4J) significantly reduced both demargination (Figure 4K) and recruitment to the heart (Figure 4L). These findings suggest that NE binding to β ARs particularly the β2 ARs are crucial in mobilizing the neutrophils from the vascular wall to the heart in response to MI. In support of this notion, we also found that β2 ARs are most abundant receptors in neutrophils (Supplemental Figure 3A).

**Figure 4.**
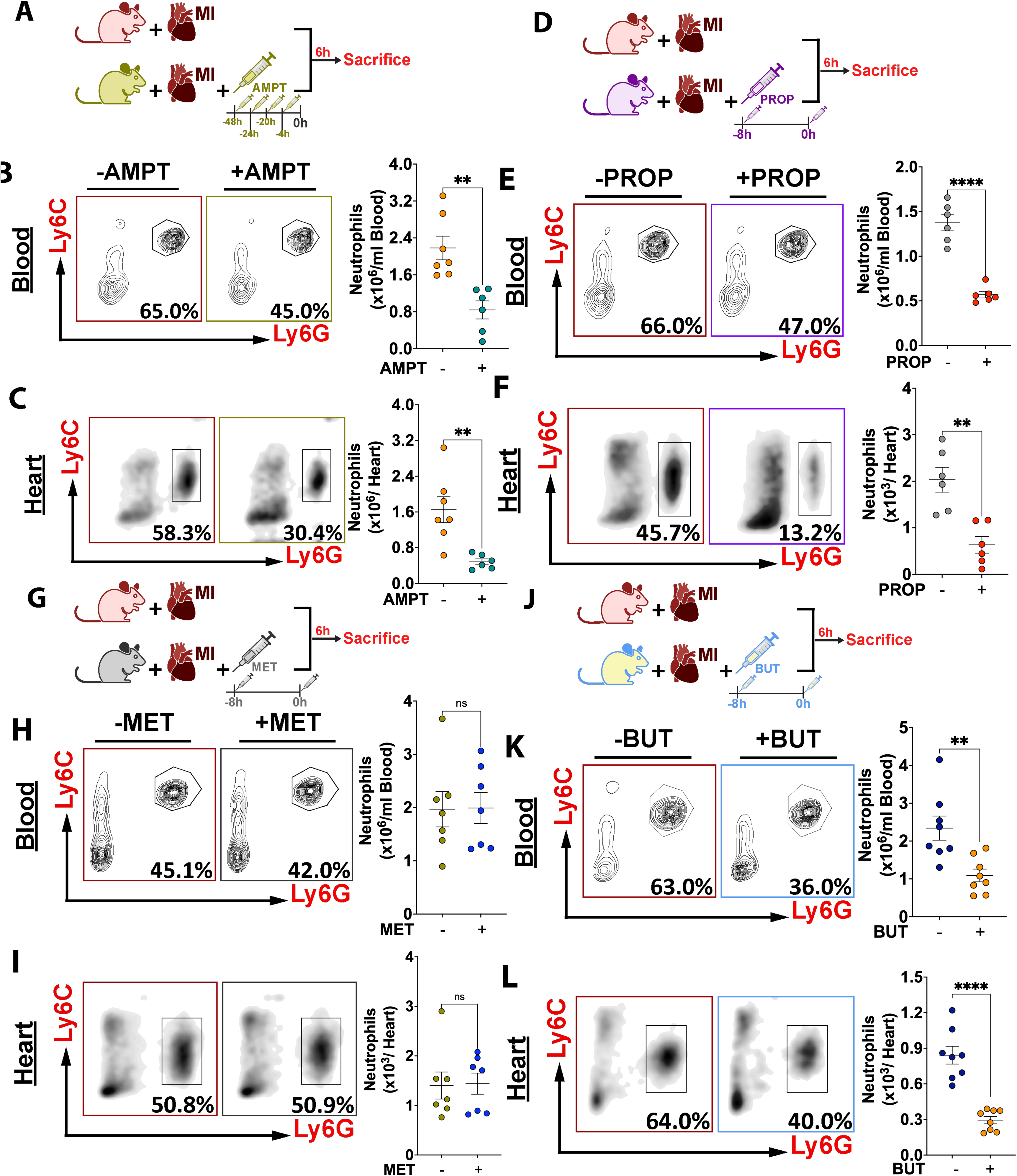
Pharmacological inhibition of β2-adrenergic receptors or NE synthesis suppresses neutrophil demargination and their recruitment to the heart. C57BL/6J WT mice were subjected to MI by permanent ligation and treated with various agents as indicated in study outlines. **A**, Schematic figure depicting study outline including administration strategy for AMPT (Alpha-Methyl-Para-Tyrosine), a NE synthesis inhibitor. **B**, Representative flow cytometric plots (left panel) and quantification (right panel) of neutrophils in the blood at 6 hours post-MI. **C**, Representative flow cytometric plots (left panel) and quantification (right panel) of neutrophils in the heart at 6 hours post-MI. **D**, Schematic figure depicting study outline to test the impact of propranolol, a non-selective β AR blocker. **E**, Representative flow cytometric plots (left panel) and quantification (right panel) of neutrophils in the blood at 6 hours post-MI. **F**, Representative flow cytometric plots (left panel) and quantification (right panel) of neutrophils in the heart at 6 hours post-MI. **G**, Schematic figure depicting study outline to test the impact of metoprolol, a selective β1 AR blocker. **H**, Representative flow cytometric plots (left panel) and quantification (right panel) of neutrophils in the blood at 6 hours post-MI. **I**, Representative flow cytometric plots (left panel) and quantification (right panel) of neutrophils in the heart at 6 hours post-MI. **J**, Schematic figure depicts study outline to test the impact of butoxamine, a selective β2 AR blocker. **K**, Representative flow cytometric plots (left panel) and quantification (right panel) of neutrophils in the blood at 6 hours post-MI. **L**, Representative flow cytometric plots (left panel) and quantification (right panel) of neutrophils in the heart at 6 hours post-MI. AMPT was administrated at 48, 24, 20, and 4 hours before MI to ensure effective suppression of NE synthesis. Propranolol (PROP), metoprolol (MET) and butoxamine (BUT) were administered 8 hours prior to surgery and once immediately after surgery. All data are means ± SEM. Statistical tests for **B** through **L** (Unpaired t test, **P<0.005, ****P<0.0001. n.s, not significant).

We next tested whether glucocorticoid stress plays any role in MI-induced neutrophil demargination. Given that DEX is used as a classical neutrophil demargination agent, we sought to clarify if blockade of glucocorticoid receptor signaling would have any impact on neutrophil demargination or their recruitment to the heart. Prior treatment of mifepristone, a potent glucocorticoid antagonist (Study outline, Supplemental Figure 3B), surprisingly had no impact on neutrophil demargination (Supplemental Figure 3C) or their recruitment to the heart (Supplemental Figure 3D) in response to MI. Together, these findings suggest that catecholamine stress, but not glucocorticoid stress promotes neutrophil demargination during the early hours after MI.

### Hematopoietic cell specific deletion of β-ARs suppresses neutrophil demargination in response to MI

Given the ability of NE in stimulating neutrophil demargination by involving one or more β-ARs, we hypothesized that myeloid cell-specific deletion of β2 or both β1and β2 (DKO) should significantly dampen demargination of neutrophils and their potential recruitment to the infarct. To test this hypothesis, we performed bone marrow transplant (BMT) studies. Briefly, we transplanted BM from WT, β2 or β1β2 AR knockout mice into age and sex matched WT mice and subjected them to MI (Study outline, Figure 5A). Following a 6-week BM reconstitution period, MI was induced, and a group of mice was euthanized within 6 hours to examine the markers of granulopoiesis, neutrophil demargination and recruitment to the heart. The remaining mice were monitored for cardiac dysfunction and survival over a period of 30 days. As expected, induction of MI significantly increased the number of neutrophils at 6 hours post-MI in the blood (Figure 5B) and hearts (Figure 5C) of recipient mice transplanted with BM from WT mice. However, deletion of β2 or both β1 and β2, significantly decreased the number of neutrophils in the blood and heart. Among the chimeric mice, mice lacking β2 ARs showed the most robust decline in neutrophil recruitment (63.5% vs 37.2%) to the heart suggesting that β2 ARs are vital for neutrophil demargination and recruitment during early hours post-MI. Interestingly, the number of neutrophils remained lower in the blood (Supplemental Figure 4A) but increased in spleens (Supplemental Figure 4B) of both chimeric mice at 30 days post-MI suggesting a sympathetic hypo-responsiveness to the loss of β AR on leukocytes^31^. However, as expected, the number of HSPCs including CMPs and GMPs in the BM did not differ either at 6 hours or 30 days post MI (Supplemental Figure 4,C and D) among the different groups. These data reiterates that early neutrophil response during MI is not driven by emergency granulopoiesis in the BM.

**Figure 5.**
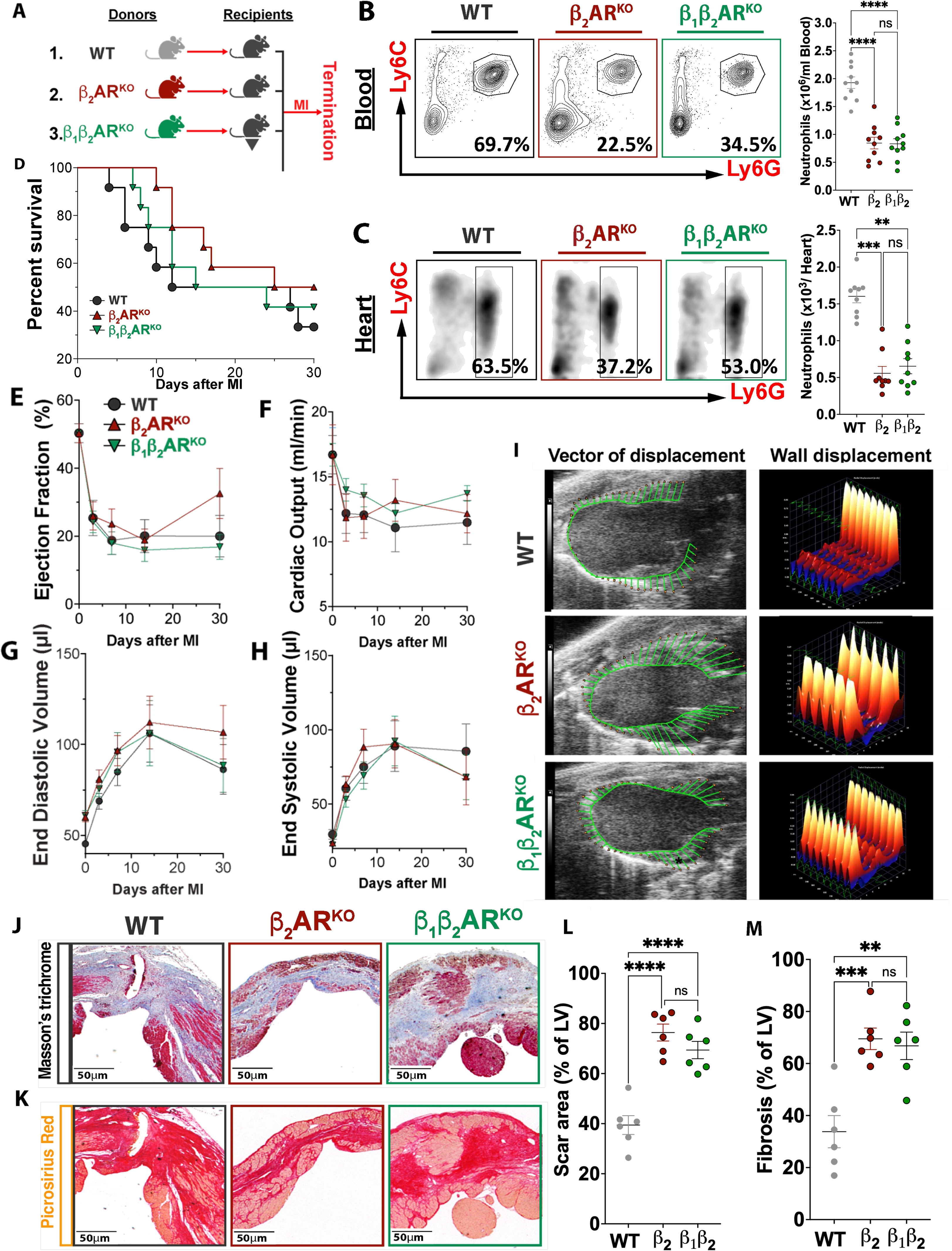
Hematopoietic cell-specific deletion of β-adrenergic receptors suppresses neutrophil demargination but does not improve survival or cardiac function following MI. **A**, Experimental overview. Bone marrow (BM) from C57BL6J WT mice, β2 or both β1 and β2 (DKO) mice were transplanted to Wild Type (WT) recipients and allowed to reconstitute for 6 weeks, after which MI was induced. **B**, Representative flow cytometric plots (left panel) and quantification (right panel) of neutrophils in the blood at 6 hours post-MI. **C**, Representative flow cytometric plots (left panel) and quantification (right panel) of neutrophils in the hearts at 6 hours post-MI. **D**, Cumulative Kaplan-Meir survival curve analysis of BMT mice monitored for 30 days post-MI. Log-rank (Mantel-Cox) test found no significant difference in survival rate among the groups. The effect of MI on cardiac function was assessed by echocardiography at day 0 (pre-MI), 3-, 7-, 14- and 30-days post-MI. Measurement of left ventricular functional parameters by echocardiography including (**E**) Ejection fraction, (**F**) Cardiac output, (**G**) End Diastolic Volume and (**H**) End Systolic Volumes. **I**, Vector diagrams showing the direction and magnitude of myocardial contraction. Three-dimensional regional wall displacement illustrations, demonstrating contraction (yellowish red) or relaxation (blue) of consecutive cardiac cycle results at 30 days post-MI. Representative histological sections (Masson’s Trichrome stain) of LV at 30 days post-MI showing (**J**) scar size and (**L**) its quantification as percentage of LV. Representative histological sections (Picrosirius stain) of LV at 30 days post-MI showing (**K**) cardiac fibrosis and (**M**) its quantification as percentage of LV. All data are means ± SEM. For cardiac function assessment, n= 5-10 mice/ group. Statistical tests for **B, L** and **M** (1-way ANOVA and Tukey’s post-hoc test, **P<0.005, ***P<0.0005, ****P<0.0001, n.s, not significant). Statistical tests for **C (**Kruskal Wallis+ Dunn’s, **P<0.005, ***P<0.0005). Statistical tests for **E** through **H**, Two-Way repeated measures ANOVA and Bonferroni multiple comparison test.

Despite a significant decrease in both neutrophil demargination and recruitment at 6 hours post-MI in β2 and DKO mice, all chimeric mice demonstrated similar survival rates (Figure 5D). Furthermore, all cardiac functional paraments tested including left ventricular ejection fraction (Figure 5E), cardiac output (Figure 5F), End Diastolic (Figure 5G) and End Systolic Volumes (Figure 5H) were similar among all 3 groups. The hearts from all groups showed a similar degree of hypokinesis (Figure 5I) although the size of scar (Figure 5J) and fibrosis (Figure 5K) was significantly increased in both β2 and DKO mice compared to WT mice. Thus, while blockade of β ARs reduces neutrophil recruitment to the heart, the lack of functional β ARs over a prolonged period post-MI does not translate into better survival rates or cardiac functional outcomes.

### Short-term, but not long-term, inhibition of β2-ARs ameliorates cardiac remodeling and improves functional parameters following MI

Previous studies have reported that depletion of neutrophils beyond the inflammatory stage (> 3 days) during MI have a negative impact on cardiac remodeling and heart failure^32^. Similarly, deletion of β ARs, particularly the β2 ARs in hematopoietic cells showed increased mortality due to impaired recruitment of leukocytes over a prolonged time following MI^31^. To examine this notion within a framework of neutrophil demargination induced by activation of β ARs during MI, we performed an *in vivo* study comparing the short-term (ST) vs long-term (LT) impact of β AR blockade using propranolol. For short-term β AR inhibition, mice were administered with one dose of propranolol 8 hours prior to and another dose immediately after surgery. Long-term (LT) inhibition involved 2 doses of propranolol every day (12 hours apart) continuously for 6 days. The control group was treated with a vehicle for 6 days (Study outline, Figure 6A). All mice were observed for 30 days while assessing cardiac function by echocardiography at day 0, day 3, day 7, day 14, and day 30.

**Figure 6.**
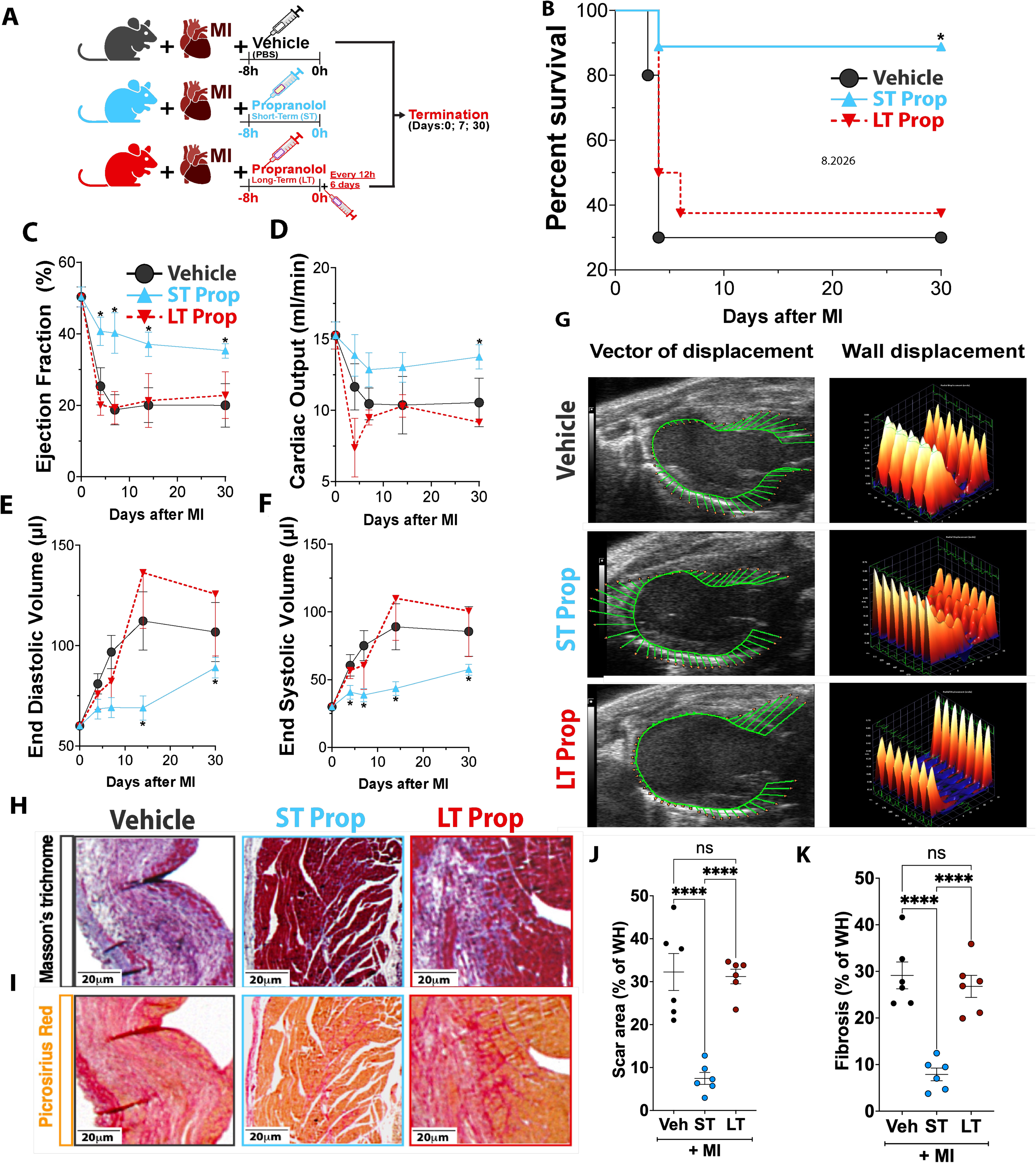
Short-term, but not long-term, inhibition of β2-adrenergic receptors improves cardiac remodeling and function following MI. **A**, Schematic figure depicting the study outline to test the effect of short term (ST) or long term (LT) blockade of β1/β2 ARs post-MI. **B**, Kaplan-Meir survival curve analysis of mice treated with or without propranolol for ST or LT. Long-rank (Mantel-Cox) test. *P <0.05 vs Vehicle and LT- Prop groups. Measurement of left ventricular functional parameters by echocardiography including (**C**) Ejection fraction, (**D**) Cardiac output, (**E**) End Diastolic Volume and (**F**) End Systolic Volumes. **G**, Vector diagrams showing the direction and magnitude of myocardial contraction. Three-dimensional regional wall displacement illustrations, demonstrating contraction (yellowish red) or relaxation (blue) of consecutive cardiac cycle results at 30 days post-MI. Representative histological sections (Masson’s Trichrome stain) of LV at 30 days post-MI showing (**H**) scar size and (**J**) its quantification as percentage of whole heart (WH). Representative histological sections (Picrosirius stain) of LV at 30 days post-MI showing (**I**) cardiac fibrosis and (**K**) its quantification as percentage of WH. All data are means ± SEM. n= 6-9 mice/ group. Statistical tests for **C** through **F** (Two-way repeated measured ANOVA and Bonferroni’s multiple comparison test, *P <0.05 vs Vehicle and LT groups at corresponding time point). Statistical tests for **J** and **K (**1-way ANOVA and Tukey’s post-hoc test, ****P <0.0001, n.s, not significant).

Contrary to the survival data in DKO mice from BMT studies, the survival rate of mice in ST group was significantly higher than those receiving LT propranolol treatment (90% vs 40%) or vehicle (90% vs.30%) (Figure 6B). This improved survival is likely due to a substantial reduction in neutrophil infiltration into the heart (Supplemental Figure 5B and 5C), which corresponded with decreased cardiac injury, as indicated by lower levels of cardiac troponin I (Supplemental Figure 5D). The mice that survived MI in ST group demonstrated a significant improvement in cardiac functional parameters including preserved ejection fraction (Figure 6C), cardiac output (Figure 6D) and lower end diastolic (Figure 6E) and systolic volumes (Figure 6F) measured at 30 days post-MI. The vector diagrams illustrating the three-dimensional regional wall displacement also supported an overall improvement in contraction and relaxation parameters (Figure 6G). The size of scar (Figure 6,H and J) as well as percentage of fibrosis (Figure 6, I and K) in ST group was also much lower than LT and vehicle groups. Additional markers including α-SMA (Supplemental Figure 5E and 5F), ERK (Supplemental Figure 5E and 5G), Col1A1 (Supplemental Figure 5I), Col3A1 (Supplemental Figure 5J), and Fibronectin I (Supplemental Figure 5K) further support the conclusion that short-term inhibition of β1/β2-adrenergic receptors significantly improves cardiac remodeling and functional recovery.

The beneficial effects of ST inhibition of β ARs with propranolol on cardiac phenotype is mainly due to the blockade of β2 ARs rather than β1 ARs. This is because, short-term inhibition of β1 AR with metoprolol (Study outline, Supplemental Figure 6A) did not improve any of the cardiac functional parameters including hypokinesia (Supplemental Figure 6B), ejection fraction (Supplemental Figure 6C), cardiac output (Supplemental Figure 6D), end diastolic (Supplemental Figure 6E) or systolic volumes (Supplemental Figure 6F) while β2 AR blockade with butoxamine significantly improved both the cardiac functional parameters (Supplemental Figure 6, B-F), scar size (Supplemental Figure 6, G and H) and fibrosis (Supplemental Figure 6, I and J). Together, these data suggest that blockade of NE binding to β2 ARs on neutrophils during the peak inflammatory stage (< 1 day) suppresses neutrophil mobilization and recruitment to the heart thus, facilitating a favorable cardiac remodeling process and better functional outcomes.

### Short-term inhibition of β1/β2-ARs using propranolol also decreases neutrophil infiltration and enhances cardiac function following ischemia-reperfusion (IR) injury

While the permanent ligation model is widely used to study ischemic injury, it does not necessarily reflect the clinical scenario of AMI, which typically involves timely reperfusion. However, since many patients still face delays in revascularization and because both permanent ligation and reperfusion injury trigger neutrophil recruitment, we aimed to determine whether ST inhibition of β1/β2 ARs in a reperfusion injury (IR) model would yield outcomes similar to those observed in the permanent ligation model. In brief, ischemia was induced by ligating the LAD artery for 50 minutes, followed by reperfusion (experimental design, Supplemental Figure 7A). One group of mice was euthanized at 6 hours post-IR to evaluate the effect of short-term propranolol (ST) treatment on cardiac neutrophil infiltration, while the remaining mice were monitored for an additional 30 days to assess long-term cardiac outcomes. As anticipated, IR injury led to a marked increase in the number of neutrophils in the heart as assessed by flow cytometry (Supplemental Figure 7B and C) and IHC (Supplemental Figure 7D and E). This was associated with elevated levels of cardiac troponin I in the plasma (Supplemental Figure 7F and G), an indicator of cardiac tissue damage. Short-term propranolol treatment significantly reduced neutrophil infiltration into the heart as well as cardiac damage as shown by reduced cTnI levels. This resulted in a significant improvement in cardiac function including all the saliant features of hypokinesia (Supplemental Figure 7H), ejection fraction (Supplemental Figure 7I), cardiac output (Supplemental Figure 7J), end systolic and diastolic volumes (Supplemental Figure 7K and L). While the hearts of vehicle treated group showed increased scar size (Supplemental Figure 7M and N) and fibrosis (Supplemental Figure 7M and O), short term blockade of β1/β2 ARs with propranolol not only prevented this but also preserved the cardiac function. These findings suggest that both ischemic and reperfusion injuries promote neutrophil mobilization to the injured heart and blocking β1/β2 ARs transiently with propranolol may be effective in preventing adverse cardiac remodeling and preserving cardiac function.

## Discussion

Myocardial infarction triggers a robust neutrophil-driven inflammatory response^4,12^. Within hours, circulating neutrophils are rapidly recruited to the infarcted tissue, where they encounter chemokines, cytokines, and DAMPs such as S100A8 and S100A9 released by necrotic cells. These signals prime the NLRP3 inflammasome and, in some neutrophils, trigger NETosis, amplifying inflammation and promoting granulopoiesis in the bone marrow (BM) via S100A8/A9 complexes^33,34^. Notably, while BM-driven granulopoiesis peaks at 24 hours post-MI, neutrophil levels in the heart and blood rise as early as 6-12 hours, indicating that a pre-existing pool of circulating or marginated neutrophils likely contributes to the initial wave.

Under homeostatic conditions, neutrophils released from the BM distribute equally between the circulating and marginated pools^35^. These compartments remain in equilibrium but are disrupted by physiological or pathological stimuli. We hypothesized that the marginated neutrophil pool serves as the primary reservoir for the early neutrophil influx post-MI. To test this, we used irradiation to deplete HSPCs in the BM and spleen. By 18 hours post-irradiation, HSPCs were eliminated, yet neutrophilia persisted following either MI or administration of DEX, suggesting demargination, rather than granulopoiesis, as the source of neutrophils. Additional experiments showed that MI exhausted the marginated pool, as DEX injection in MI mice failed to further increase neutrophil counts. Moreover, neutrophils from MI-, DEX-, or NE-treated mice shared demargination-associated features such as increased F-actin intensity, spreading, and surface CD62L expression. Importantly, BM depletion did not prevent G-CSF-induced neutrophilia post-MI, and proteomic profiling confirmed activation of pathways related to cell movement and trafficking, consistent with demargination. β-AR blockade with propranolol, particularly targeting β2-ARs, suppressed these signatures, indicating their key role in neutrophil mobilization.

To identify the mechanisms that facilitate demargination of neutrophils, we focused on two well-known stress pathways, the corticosteroid (DEX) and the catecholamine (NE) pathway. Proteomic analysis revealed minimal activation of the glucocorticoid receptor NR3C1. Furthermore, mifepristone (a glucocorticoid receptor antagonist) did not impair neutrophil recruitment post-MI, in line with reports that neutrophils express a largely inactive β isoform of the receptor^36^. These data suggest that while DEX can induce demargination experimentally, the corticosteroid pathway plays a minimal role in early neutrophil mobilization after MI. Conversely, catecholamine signaling via β-ARs appears central. MI upregulated GPCR-linked signaling molecules such as ARBK1, GRK6, and cAMP-activated protein kinase subunits KAPCA/B, which regulate actin cytoskeleton remodeling, microtubule dynamics, and cell migration^37,38^. NE administration to healthy mice mimicked the cellular responses seen post-MI, including cytoskeletal rearrangement and increased cell motility responses that were suppressed by β-AR blockade with propranolol. These findings underscore β-AR signaling, especially β2-AR, as a primary driver of early neutrophil demargination.

To further validate the role of β-ARs, we performed pharmacological and genetic interventions. AMPT (a catecholamine synthesis inhibitor), butoxamine (a β2-AR antagonist), and propranolol significantly reduced neutrophil levels in blood and heart post-MI, whereas metoprolol (β1-selective) had no effect. In BM chimeric mice, deletion of β2-AR or both β1/β2-ARs led to a sustained reduction in neutrophil burden at both acute (6 hours) and chronic (30 days) post-MI timepoints. β2-AR deletion in hematopoietic cells alone markedly reduced cardiac neutrophil infiltration, confirming its dominant role in demargination. Interestingly, double knockout (DKO) of β1/β2-ARs also reduced blood neutrophils, possibly due to impaired BM sympathetic signaling and suppression of immune-associated gene networks in the hypothalamus, as previously reported^39,40^.

Despite a significant decrease in cardiac neutrophil burden in all chimeric mice, the survival rate and the indices of cardiac function were similar to WT BMT mice. These findings are not surprising as many recent studies have suggested that depletion of neutrophils beyond the inflammatory stage (> 3 days) can cause more harm than benefit to the heart ^32,41,42^. It is important to note that all our chimeric mice were neutropenic for 30 days and thus, may have adversely impacted the resolution of inflammation ^32^. To address this, we compared short- vs long-term β2-AR inhibition using butoxamine or propranolol. Transient blockade (<12 hours) post-MI significantly reduced neutrophil burden, improved survival, and enhanced cardiac function. In contrast, prolonged inhibition or no intervention worsened outcomes. This dichotomy may stem from the dual role of neutrophils: while they promote inflammation early on, they are also essential for later healing via release of factors like S100A8/A9, which facilitate monocyte-to-macrophage conversion through NR4A1 activation. Inhibition of S100A8/A9 during early MI reduces neutrophil recruitment and improves cardiac function, supporting the idea that neutrophil-derived alarmins also aid in tissue repair.

In summary, MI induces sympathetic overdrive, triggering β2-AR-mediated neutrophil demargination and early infiltration into the heart. While essential for debris clearance, this initial neutrophil wave can exacerbate injury. Short-term β2-AR blockade attenuates early neutrophil mobilization and improves cardiac outcomes, whereas prolonged suppression may hinder resolution. These findings highlight the therapeutic potential of temporally targeted β2-AR inhibition in acute MI.

Our study has several limitations. *First*, while our data suggest systemic activation of the SNS drives demargination, we cannot rule out localized myocardial injury as a contributing factor. Notably, sham surgeries did not elicit similar neutrophil responses, supporting the systemic stress hypothesis. Second, the data from our metoprolol and the long-term β AR blockade studies including BMTs do not support the current paradigm of using β blockers in AMI patients over an extended period. These findings are in direct contrast to the current clinical guidelines that recommend the indefinite use of β blockers in AMI patients with decreased LVEF^43^. The evidence that formed the basis for the current guidelines with β-blockers was drawn from trials conducted prior to the advent of contemporary reperfusion techniques such as percutaneous coronary intervention (PCI)^44^. The general consensus was that β blockers would offer the immediate benefit of reducing the infarct size, increasing the threshold for ventricular arrhythmias and in the long run, preventing maladaptive ventricular remodeling and heart failure^45-48^. Thus, the major goal of using β blockers was to blunt sympathetic overdrive in AMI patients. However, with availability of modern revascularization strategies, anti-platelet agents, ACE inhibitors and statins, all of which are known to reduce sympathetic activation or infarct size independent of β blockade^49-51^, the benefits of using β-blockers post MI have been questioned. Although the current guidelines recommend the use of β-blockers long-term in patients with LVEF < 40%, further studies are required to determine whether short-term inhibition of β2 AR with propranolol or carvedilol is superior to the indefinite use of these agents particularly in patients who are at low risk for cardiogenic shock.

## Supplemental Material

Supplemental materials and methods Supplementary Figures. 1-7 Supplemental Figure legends

## Acknowledgements

The authors thank the staff of flow cytometry, microscopy, histology cores at OSU and OUHSC for their assistance. This work was supported by funds from the NIH (HL137799, HL156856) and AHA (TPA97002) to PN. DT was supported by NIH (HL147841).P.N. and A.D., conceived and designed the study, analyzed the data, created figures, and wrote the main manuscript that was revised and approved by all authors. A.D., M.M., K.P., R.J., N.N., and W.Q. performed animal experiments, surgeries, BMTs, FACS, mouse echocardiography, proteomics, microscopy, and histological analysis. T.D., S.S., D.T., and A.M contributed to manuscript writing, editing, discussion and critical inputs. V.I.S., E.U., R.M.B. and F.A. performed proteomics analysis and contributed to writing and editing of manuscript.

