## Supplemental Document for "β2 adrenergic receptors orchestrate neutrophil demargination and recruitment to the ischemic heart following myocardial infarction"

**Institutions**

^1^ Department of Internal Medicine, Section of Cardiovascular Diseases, College of Medicine, University of Oklahoma Health Sciences Center, Oklahoma City, OK, USA.

^2^ Department of Surgery, Division of Cardiac Surgery, Ohio State University Wexner Medical Center, Columbus, OH, USA

^3^ Proteomics Department, Institute of Cellular Biology and Pathology “Nicolae Simionescu” of the Romanian Academy, Bucharest, Romania.

^4^ Aging+ Cardiovascular Discover Center, Lewis Katz School of Medicine, Temple University, Philadelphia, USA.

^5^ Baker Heart and Diabetes Institute, Division of Immunometabolism, Melbourne, Australia.

**Short title:** Neutrophil β2 receptors promote demargination.

***Corresponding author,**

Prof. Prabha Nagareddy, PhD, FAHA

1122 NE, 13th Street, Suite 333B

Oklahoma City OK 73117, USA

**Materials and Method details.**

**Mice and Treatments:** Male mice aged between 10-12 weeks including Wild Type (C57BL/6), β2AR^-/-^ and β1/β2 AR^-/-^ (DKO)*,* in C57BL/6 backgrounds were used for this study. Since MI is known to impact cardiac remodelling differently in males and females, we used only male mice to reduce biological variation and prevent the confounding effects of sex on data interpretation, particularly that related to neutrophil behaviour and its impact on cardiac remodelling post-MI. Wild Type mice were purchased from the Jackson Laboratory while β2AR^-/-^ and β1/β2AR^-/-^ (DKO) mice were provided by Dr. Doug Tilley, Temple University, Philadelphia. All mice were maintained in pathogen-free environments at The Ohio State University (OSU) and University of Oklahoma Health Sciences Center (OUHSC) animal facilities. Procedures were conducted under approval from the OUHSC and OSU’s Institutional Animal Care and Use Committee, in accordance with the NIH Guide for the Care and Use of Laboratory Animals. All mice were fed with a normal chow diet *ad libitum* and randomly assigned to experimental groups.

**Drugs and Administration:** All drug solutions meant for *in vivo* studies were prepared under sterile conditions according to manufacturer’s instructions and administered according to approved IACUC protocols.

Propranolol HCl (30mg/kg, i.p., Millipore-Sigma P0884)

Butoxamine HCl (12.5mg/kg, i.p., Sigma-Aldrich B1385)

Metoprolol Tartrate (12.5mg/kp, i.p., Millipore-Sigma M5391)

Alpha-Methyl-DL-Tyrosine, AMPT (40mg/kg, s.c., Millipore-Sigma 120693)

Dexamethasone (5mg/kg, i.v, Millipore-Sigma D2915)

Granulocyte Colony-Stimulating Factor, G-CSF (100mg/kg, i.p., Stem Cell Tech 78014.2)

Norepinephrine HCl (2mg/kg, i.p., Millipore-Sigma A7256)

Mifeprestone (30mg/kg., s.c., Millipore-Sigma M8046)

ADM3100 (10mg/kg, s.c., Millipore-Sigma A5602)

5-Ethynl-2’-Deoxyuridine, EdU (16mg/kg, i.p., Thermo Fisher C10340)

All other materials and reagents are listed in Supplemental Table -S1

**Method Details**

**Mouse Model of Myocardial Infarction**: Myocardial infarction (MI) was induced by permanent ligation of the left anterior descending artery. Twenty-four to 48 hours before surgery, the fur on ventral surface in and around left chest area was cleared off using depilatory cream. Briefly, the mice were anesthetized with 2% isoflurane anaesthesia using a continuous isoflurane delivery system and with an endotracheal tube respiration was controlled with use of a rodent ventilator. A small longitudinal incision (≤5 mm) was made in the centre of the throat to locate the trachea for intubation. After confirming the intubation, 4-0 vicryl sutures were used to close the skin in the throat. The line between left pectoralis and major muscles was identified and then an oblique incision (~7mm) was made to expose the heart and left coronary artery. An 8-0 suture (silk or nylon) was placed around the coronary artery and ligated to create the ischemia. Animals were monitored for the depth of anaesthesia at least every 10 to 15 minutes throughout the surgery. After controlling the exhaust pressure to restore the lungs to normal, 5-0 vicryl sutures were used to close the muscle layer while 4-0 vicryl sutures were then to close the skin on the chest. The animals were gradually weaned off the ventilator during recovery from anaesthesia and monitored further to confirm that they have resumed spontaneous respiration in a normal rhythm without assistance during the postoperative period. The sham group underwent the same surgical procedure, except that the ligature was passed under the LAD but not tied. Immediately after surgery, all mice were injected with a long-acting analgesic, buprenorphine. No-MI mice did not undergo either LAD or sham surgery.

**Mouse Model of Myocardial Ischemia-Reperfusion Injury (IRI)**: Myocardial ischemia-reperfusion injury was induced in adult male C57BL/6J mice (10–12 weeks old) as previously described, with minor modifications. Briefly, mice were anesthetized with isoflurane (1–2% in oxygen) and placed on a heating pad to maintain body temperature. A left thoracotomy was performed to expose the heart, and a 7-0 silk suture was placed around the left anterior descending (LAD) coronary artery approximately 1–2 mm below the tip of the left atrium. To induce ischemia, the suture was tightened and secured with a small piece of PE-10 tubing to occlude the LAD for 50 minutes. Ischemia was confirmed by pallor of the left ventricle and ECG changes. After 50 minutes, the ligature was released to allow reperfusion, verified by return of myocardial color. Mice were monitored during the reperfusion period and euthanized at indicated time points for tissue and plasma collection. The sham group underwent the same surgical procedure, except that the ligature was passed under the LAD but not tied. Immediately after surgery, all mice were injected with a long-acting analgesic, buprenorphine. No-MI mice did not undergo either ligation or sham surgery.

**Transthoracic Echocardiography:** Echocardiography was performed using the Vevo 3100 high-resolution ultrasound imaging system with a MX400 transducer (Visual Sonics Inc., Toronto CA). Mice were sedated with 2% isoflurane and placed on temperature-maintained platform in a supine position. Hair was removed from the chest using depilatory cream, warmed ultrasound transmission gel placed on the chest and left parasternal long axis view and basal short axis views acquired. Ejection fraction, stroke volume and cardiac output were evaluated on the left parasternal long axis view while fractional shortening was calculated from the short axis view using the Vevo 3100 software. Core body temperature was maintained at 37°C using a heated platform and infrared heat lamp throughout the procedure. Mice with heart rates below 400 bpm (beats per minute) were excluded from the analysis.

**Bone Marrow Transplants (BMT):** WT mice were lethally irradiated (with 13 Gy from a Cesium gamma source in 2 doses separated by 4 hours) and transplanted with BM (5x10^6^ cells) from donor mice. Chimerism was confirmed using either flow cytometry or qRT-PCR analysis of respective proteins/ genes (mRNA) in peripheral blood cells at ~6 weeks after transplantation. Once the chimerism was confirmed, MI was induced as described above.

**White Blood Cell (WBC) Counts:** Total WBC in freshly collected mouse blood was measured using hematology analyser (Element HT-5, Heska, Inc.,) or a K2 Fluorescent Viability Cell Counter (Nexcelom Biosciences, MA).

**Flow Cytometry Analysis.**

**Blood and Bone Marrow Leukocytes:** Briefly, fresh blood from mice was collected via tail bleeding directly into microcentrifuge tubes containing EDTA (5 mM) using heparin coated capillary tubes and immediately placed on ice. A small amount of blood was set aside for separation of plasma and for analysis of complete blood cell count (CBC) while the remaining blood was transferred to 15 ml centrifuge tubes and red blood cells were lysed using BD Pharm Lyse (BD Biosciences, San Jose, CA), washed, rinsed and resuspended in PBS or FACS buffer (Hanks balanced salt solution, HBSS + 0.1% BSA w/v, 5mM EDTA) for staining.

For isolation of BM, femurs and tibias from each mouse were isolated, cleaned and their ends were cut open with a sharp scissor to expose lumen. Bone marrow was flushed with ice cold PBS using a 23-gauge, 10 ml syringe directly through a 40 mM nylon mesh strainer on a 50 ml centrifuge tube. The BM samples were next spun at 300g for 10 min at 4ºC. Following aspiration of supernatant, red blood cells were lysed using RBC lysis buffer, washed twice with PBS, and immediately stained with a fixable Live/Dead marker (1 ml per million cells in 1 ml for 30 minutes, Molecular Probes, Eugene OR). The cells were then washed and resuspended in 100 ml of FACS buffer containing a cocktail of antibodies directed against different types of leukocytes. In experiments that required intracellular staining, the surface labelled cells were fixed and permeabilized with fix/ perm buffer (30 minutes) and then stained with intracellular antibodies (e.g., S100A9, IL1b) in perm/ wash buffer. The stained cells were analysed on a LSRII flow cytometer using FACS DiVa software or Cytek Northern Lights flow cytometer (Cytek). Once the debris, doublets (by FSC-H vs. FSC-A) and dead cells, leukocytes were identified as described previously^12,13^.

**Heart Leukocytes:** For flow cytometric analysis of leukocytes in the heart, a single cell suspension was first prepared. Briefly, hearts were perfused with ice cold PBS to remove blood cells, harvested, cleared of adhering tissues and atria, minced into 2-3 large pieces, transferred to GentleMACS C tubes (Miltenyi Biotec) containing either collagenase type II (1mg/ml) alone or a cocktail of collagenase I (450U/ml), collagenase XI (125U/ml), hyaluronidase type I-s (60U/ml) and DNase (60U/ml) and processed into a fine suspension using a tissue dissociator (GentleMACS Octo Dissociator, Miltenyi Biotec). Digestion was enabled by alternating between dissociation and incubation in a thermomixer (37ºC, 15 minutes each time) with gentle agitation. The single cell suspension was then filtered through a 100 μM strainer into a 50 ml tube, rinsed with FACS buffer, centrifuged at 300g for 10 min at 4ºC. The resultant pellet was washed, resuspended in 1 ml FACS buffer to determine cell count (K2 Cell Counter, Nexcelom Biosciences) followed by staining for Live/Dead marker, surface and intracellular proteins as described above. Data were acquired on Cytek Northern lights or LSR Fortessa flow cytometer (BD Biosciences, San Jose CA) using FACS Diva software, and analysis was performed with FlowJo software (Ashland, OR). Once the doublets (by FSC-H vs. FSC-A) and dead cells (Live vs. Dead) were excluded, neutrophils were identified as CD45^+^, Ly6G^hi^ cells and monocytes as CD45^+^, Ly6G^-^, F4/80^-^, CD11b^+^, CD115^+^ and further classified as Ly6-C^hi^ and Ly6-C^lo^ cells.

**Hematopoietic Stem Cells in the BM:** Hematopoietic stem and progenitor cells from the BM were prepared as described above. After cell count and staining for Live/Dead markers, the cells were resuspended in a cocktail of antibodies first against lineage-committed cells (B220, CD19, CD11b, CD3e, TER-119, CD2, CD8, CD4, Ly6-C/G) followed by markers of stem cells, proliferation (e.g., Ki67) and EdU. Hematopoietic stem and progenitor cells were first identified as Lin^-^, Sca1^+^ and ckit^+^ (LSK) while the hematopoietic progenitor subsets were separated by using antibodies to CD16/CD32 (FcgRII/III) and CD34. Common Myeloid Progenitors (CMP) were identified as Lin^-^, Sca1^-^, ckit^+^, CD34^int^, FcgRII/III^int^, Granulocyte Macrophage Progenitors (GMP) as Lin^-^, Sca1^-^, ckit^+^, CD34^int^, FcgRII/III^hi^. Common lymphoid progenitors (CLP) were identified as Lin^-^, Sca1^lo^, ckit^lo^ and IL7R+ cells. Hematopoietic stem cells (HSCs) were identified as Lin^-^, Sca1^+^, ckit^+^, CD48^-^ and CD150^+^. Cellular proliferation in hematopoietic stem and progenitor cells was assessed by measuring the incorporation of EdU and /or Ki67 according to the manufacturer’s protocol (Click-iT™ Plus EdU, Molecular Probes, Eugene OR). Flow cytometry was performed using an LSR Fortessa or Cytek Northern Lights (Cytek) using FACS DiVa or SpectroFlo software. All flow cytometry data were analyzed using FlowJo software.

**Confocal Microscopy:** Neutrophils from whole blood were sorted using Neutrophil Isolation kit (130-097-658 Miltenyi Biotec), neutrophils were then permeabilized with BD Cytoperm/Cytofix (BD554722), Fc receptors blocked with Fc Block (TruStain FcX PLUS anti-mouse CD16/CD32, 156603, Biolegend) and stained with Phalloidin, Fluroescein isothiocyanate labelled (P5282, Millipore Sigma) for 60mn and mounted using ProLong Gold antifade reagent with DAPI (P36935, Invitrogen). Neutrophil purification efficiency was assessed by flow cytometry (CD45, CD11b, Ly6G). Confocal images were obtained using Olympus FV3000 Multi confocal.

**Histology of Heart Sections**: The hearts were harvested in diastole with saturated KCl and CdCl (100 mM) injected through the apex into the left ventricular (LV) cavity. The LV apex was then cannulated, and the heart perfused with PBS, followed by 10% buffered formalin at 75 mmHg, with the severed inferior vena cava serving as the outlet. The hearts were then cut into 2-mm cross-sectional slices and processed for paraffin embedding. The slices were cut into 4 μ M sections for histologic examination and picrosirius staining. Images were acquired digitally, and the areas measured using NIH Image J software.

**Proteomics**

**Cell Lysis:** Cell pellets were washed with PBS three times before resuspend in 100 µL of 5% SDS buffer in 50 mM TEAB (Triethylammonium bicarbonate) solution. Samples then were vortexed briefly before proceeding to sonication by using the Bioruptor® Pico (Diagenode, Denville, NJ) following manufacturer’s suggested protocol. Briefly, sonication temperature was set at 4C; sonication cycle was set at 30 sec on and 30 sec off. A total of 10 cycles were done for the cell pellets. (Sometimes, DNAs are not sheared well in SDS buffer so a probe homogenizer can be used to shear DNAs in that case). After sonication, samples were centrifuged at 14,000 rpm for 15 min at 4°C to remove any remaining insoluble material. Concentration of the proteins were measured using Qubit fluorometer (ThermoFisher Scientific).

**Protein Digestion via Strap:** 50 µg of the samples were taken for trypsin digestion. 5 µL of 50 mM ABC containing 5 µg/µL DTT was added, and the sample was incubated at 65^0^C for 15 min followed by addition of 5 µL of 50 mM ABC containing 15 µg/µL iodoacetamide (incubated at RT for 15 min in the dark). Samples were then acidified by adding 12% phosphoric acid (1:10 v/v acid to sample). For every 25 µL of samples, 165 µL of TEAB (Triethylammonium bicarbonate, 1M)/MeOH (10:90 v/v) was added and then loaded to S-trap for further washes. Samples was centrifuged at 4000g for 3 min (4C) to remove supernatant. 150 µL of TEAB (Triethylamonium bicarbonate, 1M)/MeOH (10:90 v/v) was added to the trap as wash solution and the trap was washed 3-6 times depends on the initial loading volume. After the final wash, sequencing grade trypsin dissolved in 50 mM TEAB was added and digested O/N at 37^0^C. The following day, peptides were eluted from the trap by adding 40 µL of 50mM TEAB, 0.1% FA and 0.1% FA in Acetonitrile (50:50), sequentially. The sample was pooled together and dried in a vacufuge and resuspended in 20 µL of 50 mM acetic acid. Peptide concentration was determined by nanodrop (A280 nM).

**Orbitrap Fusion:** Capillary-liquid chromatography-nanospray tandem mass spectrometry (Capillary-LC/MS/MS) of protein identification was performed on a Thermo-Scientific orbitrap Fusion mass spectrometer equipped with an EASY-Spray™ Sources operated in positive ion mode. Samples (6.4µL) were separated on an easy spray nano column (PepmapTM RSLC, C18 3µ 100A, 75µm X150mm Thermo Scientific) using a 2D RSLC HPLC system from Thermo Scientific. Each sample was injected into the µ-Precolumn Cartridge (Thermo Scientific) and desalted with 0.1% Formic Acid in water for 5 minutes. The injector port was then switched to inject, and the peptides were eluted off the trap onto the column. Mobile phase A was 0.1% Formic Acid in water and acetonitrile (with 0.1% formic acid) was used as mobile phase B. Flow rate was set at 300 nL/min. Mobile phase B was increased from 2% to 20% in 105 min and then increased from 20-32% in 10 min and again from 32-95% in 1 min and then kept at 95% for another 4 min before being brought back quickly to 2% in 1 min. The column was equilibrated at 2% of mobile phase B (or 98% A) for 15 min before the next sample injection.

MS/MS data was acquired with a spray voltage of 1.95 KV and a capillary temperature of 305 °C is used. The scan sequence of the mass spectrometer was based on the preview mode data dependent TopSpeed™ method: the analysis was programmed for a full scan recorded between m/z 375-1500 and a MS/MS scan to generate product ion spectra to determine amino acid sequence in consecutive scans starting from the most abundant peaks in the spectrum in the next 3 seconds. To achieve high mass accuracy MS determination, the full scan was performed at FT mode and the resolution was set at 120,000 with internal mass calibration. Three compensation voltage (cv=-50, -65 and -80v) were used for samples acquisition. The AGC Target ion number for FT full scan was set at 4 x 105 ions, maximum ion injection time was set at 50 ms and micro scan number was set at 1. MSn was performed using HCD in ion trap mode to ensure the highest signal intensity of MSn spectra. The HCD collision energy was set at 32%. The AGC Target ion number for ion trap MSn scan was set at 3.0E4 ions, maximum ion injection time was set at 35 ms and micro scan number was set at 1. Dynamic exclusion is enabled with a repeat count of 1 within 60s and a low mass width and high mass width of 10ppm.

The raw files were searched using Mascot Daemon by Matrix Science version 2.5.1 (Matrix Science, Boston, MA) via ProteomeDiscoverer (Thermo) against the database provided by the user. The mass accuracy of the precursor ions was set to 10 ppm, accidental pick of 113C peaks was also included into the search. The fragment mass tolerance was set to 0.5 Da. Carbamidomethylation (Cys) is used as a fixed modification and considered variable modifications were oxidation (Met) and deamidation (N and Q). Four missed cleavages for the enzyme were permitted. A decoy database was also searched to determine the false discovery rate (FDR) and peptides were filtered according at 1% FDR. Proteins identified with at least two unique peptides were considered as reliable identification. Any modified peptides are manually checked for validation.

**Quantitation:** Label free quantitation was performed using the spectral count approach, in which the relative protein quantitation is measured by comparing the number of MS/MS spectra identified from the same protein in each of the multiple LC/MSMS datasets. Scaffold (Proteome Software, Portland, OR) was used for data analysis. Student-t test was performed by scaffold to evaluate if the folder change for certain proteins is significant (p<0.05).

**Protein inference and bioinformatic analysis:** The resulted MS raw data were processed by Proteome Discoverer V2.4 (Thermo Fisher Scientific), using the UniProtKB/Swiss-Prot Mus musculus reference protein database for protein identification. Search parameters included: enzyme: trypsin; maximum miss cleavages: 2; fixed modification: cysteine carbamidomethylation; variable modifications: oxidation of methionine and deamidation of asparagine and glutamine. The criteria for protein identification also contained the detection of at least 2 unique peptide per protein and the protein false discovery rate (FDR) target to be below 0.05. Label-free relative protein quantitation on the precursor level was performed using the same software platform, which aligns chromatograms, extracts ion peaks and integrates them to compare peptide/protein spectral abundance along the different datasets. We enabled 90% of replicate features and the ANOVA hypothesis test to determine the statistical significance associated with the protein ratio calculation. Normalization was done using the total peptide amount on controls average.

The differentially abundant proteins (DAPs) were identified with an abundance ratio ≥ 1 and an adjusted P-value ≤ 0.05, based on the Benjamini-Hochberg algorithm, was set as the cut-off criterion.

**Measurement of Cardiac Troponin I (cTnI) Levels**

Plasma samples were collected from mice at designated time points following myocardial infarction (MI) or IR, with or without propranolol treatment. Cardiac troponin I levels were quantified using the Mouse Cardiac Troponin I (TNNI3) ELISA Kit (Thermo Fisher Scientific, Cat# EEL112), following the manufacturer’s protocol. Concentrations were determined using a standard curve generated from known calibrators provided with the kit.

**Supplemental Table S1: Key Resources and Reagents**

| **REAGENT or RESOURCE** | **SOURCE** | **IDENTIFIER** |
| --- | --- | --- |
| **Antibodies** | | |
| Anti-mouse CD45 (clone 30-F11) | BioLegend | 103116 |
| Anti-mouse CD11b (clone M1/70) | BioLegend | 101228 |
| Anti-mouse CD115 (clone AFS98) | BioLegend | 135515 |
| Anti-mouse Ly6-G (clone 1A8) | BioLegend | 127612 |
| Anti-mouse Ly6-C (clone HK1.4) | BioLegend | 128018 |
| Anti-mouse Gr1 (clone RB6-8C5) | ThermoFisher | 108428 |
| Anti-mouse B220 (clone RA3-6B2) | ThermoFisher | 11-0452-85 |
| Anti-mouse CD11b (clone M1/70) | BD Biosciences | 553310 |
| Anti-mouse CD19 (clone 1D3/CD19) | BioLegend | 152404 |
| Anti-mouse CD3e (clone 145-2C11) | ThermoFisher | 11-0031-85 |
| Anti-mouse TER-119 (clone TER-119) | ThermoFisher | 11-5921-85 |
| Anti-mouse CD2 (clone RM2-5) | ThermoFisher | 11-0021-85 |
| Anti-mouse CD8a (clone 53-6.7) | ThermoFisher | 11-0081-85 |
| Anti-mouse CD4 (clone GK1.5) | ThermoFisher | 11-0041-85 |
| Anti-mouse Gr1 (clone RB6-8C5) | ThermoFisher | 11-5931-85 |
| Anti-mouse CD150 (clone TC15-12F12.2) | BioLegend | 115922 |
| Anti-mouse CD34 (clone 581) | BioLegend | 343512 |
| Anti-mouse cKit (clone 2B8) | BioLegend | 105835 |
| Anti-mouse CD16/32 (clone 93) | BioLegend | 101308 |
| Anti-mouse Sca1 (clone D7) | BioLegend | 108114 |
| Anti-mouse CD48 (clone HM48-1) | BioLegend | 103432 |
| Anti-mouse CD45.2 (clone 104) | ThermoFisher | 25-0454-82 |
| Anti-mouse CD45.1 (clone A20) | ThermoFisher | 12-0453-82 |
| Anti-mouse Annexin V | BioLegend | 640943 |
| Anti-mouse Ki67 (clone SolA15) | ThermoFisher | 12-5698-82 |
| **Chemicals, Peptides, and Recombinant Proteins** | | |
| Fixable Live/Dead marker | ThermoFisher | C2926 |
| LPS from E. coli | Enzo Life Sciences | ALX-581-012 |
| Acridine orange & Propidium Iodide) | Nexcelom | tlrl-vad |
| Weigert’s iron hematoxylin solution A | EMS | 26758-01 |
| Weigert’s iron hematoxylin solution B | EMS | 26758-02 |
| Collagenase I | Sigma-Aldrich | C0130 |
| Collagenase XI | Sigma-Aldrich | C7657 |
| DNAse I | Sigma-Aldrich | D5319 |
| Hyaluronidase | Sigma-Aldrich | H3506 |
| DAPI | ThermoFisher | F10347 |
| TNF-α | Sigma-Aldrich | T7539 |
| **Critical Commercial Assays** | | |
| Mouse Neutrophil Isolation kit | Miltenyi Biotec | 130-097-658 |
| BD Pharm Lyse | BD Biosciences | 555899 |
| Click-iT™ Plus EdU Kit | ThermoFisher | C10640 |
| Ambion Purelink RNA Mini Kit | ThermoFisher | 12183025 |
| SuperScript IV VILO Master Mix | ThermoFisher | 11766050 |
| Fast SYBR Green Master Mix | ThermoFisher | 4385612 |
| Total RNA Isolation Kit | ThermoFisher | AM1931 |
| BCA Protein Assay kit | ThermoFisher | 23225 |
| **Animals** | | |
| C57BL/6J | JAX Laboratories | 000664 |
| **Software** | | |
| Vevo 3100 software | VisualSonics Inc | <https://www.visualsonics.com> |
| FACS DiVa | BD Biosciences | <http://www.bdbiosciences.com> |
| FlowJo software (Ashland, OR) | FlowJo | <http://www.flowjo.com> |
| NIH ImageJ | NIH | <http://imagej.net> |
| Gen5 | BioTek | <https://www.biotek.com> |
| Spectraflo | Cytek | https://cytekbio.com/ |
| **Equipment/ Instruments** | | |
| Vevo 3100 High-resolution US imaging system | VisualSonics Inc., | https://www.visualsonics.com |
| K2 Fluorescent Viability Cell Counter | Nexcelom | https://www.nexcelom.com |
| LSRFortessa | BD Biosciences |  |
| LSRII Flow Cytometer | BD Biosciences | https://www.bdbiosciences.com/en-in/instruments/research-instruments/research-cell-analyzers/lsrfortessa |
| Cytek Northern Lights | Cytek | https://cytekbio.com/pages/northern-lights |
| GentleMACS Octo Dissociator | Miltenyi Biotec | https://www.miltenyibiotec.com |
| Element HT5 | Heska Inc., | https://www.heska.com/product/element-ht5/ |

**Supplemental Figure Legends**

**Supplemental Figure 1. Demargination is the predominant source of neutrophils to the ischemic heart during the early hours after MI.**

**A,** Representative flow cytometric plots (top panel) and quantification (bottom panel) of neutrophils in the blood over a period of 24 hours after administration of either nor-epinephrine (NE) or dexamethasone (DEX) to male C57BL/6 WT mice. The impact of total body irradiation (TBI) on HSCs, MPCs, CMPs and GMPs in the BM (**B**) and spleen (**C**) at various time (hours) points as indicated in individual figures. The impact of total body irradiation (TBI) on neutrophils and total live cells in the blood (**D**), BM (**E**) and Spleen (**F**) at various time points as indicated in the figures. **G**, A schematic figure depicting the experimental strategy to study the impact of DEX (administered at 6 hours post-MI) on neutrophil demargination and infiltration to the heart. **H**, Representative flow cytometric plots (left panel) and quantification (right panel) of neutrophils in the blood at 6 hours after DEX administration. **I**, Representative flow cytometric plots (left panel) and quantification (right panel) of neutrophils in the heart at 6 hours after DEX administration. **J**, A schematic figure depicting the experimental strategy to study the impact of G-CSF administration (given at 6 hours post-MI) on neutrophil number in the BM, blood and heart post-MI. **K**, Representative flow cytometric plots (left panel) and quantification (right panel) of neutrophils in the BM at 6 hours after G-CSF administration. **L**, Representative flow cytometric plots (left panel) and quantification (right panel) of neutrophils in the blood at 6 hours after G-CSF administration. **M**, Representative flow cytometric plots (left panel) and quantification (right panel) of neutrophils in the heart at 6 hours after G-CSF administration. The number in the parentheses in figures **1H**, **I**, **K**, **L** and **M** indicate the percentage of neutrophils (% CD45). All data are means ± SEM. n= 5-6 mice/ group. Statistical tests for **A**, **D, E** and **F**, Unpaired t test, ^X, *, ^, V^ P <0.05 vs corresponding baseline for each cell type. Statistical tests for **B** and **C**, 1-way ANOVA and Tukey’s post-hoc test, ^X, *, ^, V^ P <0.05 vs corresponding baseline for each cell type. Statistical tests for **H, I, K, L** and **M,** (Unpaired t test, *P <0.05, n.s, not significant). All cells were gated after eliminating debris, dead and clustered cells. Neutrophils were gated from CD45^+^ cells. Hematopoietic stem and progenitor cells (HSPCs) were first identified as Lin^-^, Sca1^+^ and ckit^+^ (LSK/ MPCs, Myeloid Progenitor Cells) while the hematopoietic progenitor subsets were separated further by using antibodies to CD16/CD32 (FcγRII/III) and CD34. Common Myeloid Progenitors (CMP) were identified as Lin^-^, Sca1^-^, ckit^+^, CD34^int^, FcγRII/III^int^, Granulocyte Macrophage Progenitors (GMP) as Lin^-^, Sca1^-^, ckit^+^, CD34^int^, FcγRII/III^hi^. Hematopoietic stem cells (HSCs) were identified as Lin^-^, Sca1^+^, ckit^+^, CD48^-^ and CD150^+^. MI, Myocardial infarction; TBI, Total body irradiation; DEX, Dexamethasone; NE, Nor-epinephrine; G-CSF, Granulocyte Colony Stimulating Factor and BM, Bone Marrow.

**Supplemental Figure 2. Proteomic analysis of pathways representing actin cytoskeleton remodelling, glucocorticoid receptor and GPCR downstream signalling in demarginated neutrophils following MI.**

MI was induced in healthy WT C57BL/6J mice and at 6 hours post-MI, neutrophils from peripheral blood were sorted using magnetic beads and subjected to whole cell proteomics analysis**.**

**A**, Principal component analysis (PCA) plot depicting the quantitative separation of the proteomes associated with the MI group (blue) and the Sham group (orange). Histograms representing gene ontology (biological process, cellular component and molecular functions**, B-D**) as well as pathway enrichment (KEGG, Reactome and Wiki Pathways, **E-G**) analyses of the differentially abundant proteins (DAPs) in MI vs Sham groups. The analysis is based on the statistical over-representation of DAPs using the STRING (Search Tool for Retrieval of Interacting Genes/ Proteins) bioinformatics tool. A higher -log10 (FDR) value indicates greater significance of the enrichment. **H**, Univariate, significance (P <0.05) vs. fold change analyses highlighting several significantly deregulated proteins representing GPCR downstream signalling, Actin cytoskeleton remodelling and glucocorticoid receptor signalling. Visualized by a volcano plot to demonstrate the level of significance and magnitude of changes observed in the proteomic data. MI, Myocardial infarction, GPCR, G-Protein Coupled Receptor.

**Supplemental Figure 3. Adrenergic receptor expression in neutrophils and the impact of mifepristone, a glucocorticoid receptor antagonist, on neutrophil demargination and recruitment to the heart following MI.**

**A,** Relative abundance of mRNA levels of various adrenergic receptors in neutrophils sorted from the blood and BM of healthy WT mice. **B**, Schematic figure depicting the experimental strategy to study the impact of mifepristone on neutrophil demargination. **C**, Representative flow cytometric plots (top panel) and its quantification (bottom panel) of neutrophils in the blood at 6 hours post-MI. **D**, Representative flow cytometric plots (top panel) and its quantification (bottom panel) of neutrophils in the heart at 6 hours post-MI. All data are means ± SEM. n= 4-7 mice/ group. Statistical tests for **A** (P <0.05 vs all other groups, 1-way ANOVA and Tukey’s post-hoc test), **B** and **C,** Unpaired t test, n.s, not significant. *Adrb1, Adrb2, Adrb3, Adra1* and *Adra2* represent adrenergic β1, β2, β3, α1 and α2 receptors respectively. MI, Myocardial infarction; MFP, Mifepristone.

**Supplemental Figure 4. Effect of hematopoietic cell-specific deletion of β-adrenergic receptors on neutrophil numbers and HSPCs in the BM following MI.**

Bone marrow (BM) from male C57BL6 WT mice, β2 or both β1 and β2 (double knockout) mice in BL6 background were transplanted to male Wild Type (WT) recipients and allowed to reconstitute for 6 weeks, after which MI was induced. Quantification of neutrophils in the blood (**A**) and spleen (**B**) of WT recipients at 30 days post-MI. **C**, Quantification of CMPs in the BM at 6 hours (top panel) and 30 days (bottom panel) post-MI. **D**, Quantification of GMPs in the BM at 6 hours (top panel) and 30 days (bottom panel) post-MI. Statistical tests for **A** through **D**, 1-way ANOVA and Tukey’s post-hoc test, ^*^P <0.05 vs all other groups. All cells were gated after eliminating debris, dead and clustered cells. Neutrophils were gated from CD45^+^ cells. Hematopoietic stem and progenitor cells (HSPCs) were first identified as Lin^-^, Sca1^+^ and ckit^+^ (LSK) while the hematopoietic progenitor subsets were separated further by using antibodies to CD16/CD32 (FcγRII/III) and CD34. Common Myeloid Progenitors (CMP) were identified as Lin^-^, Sca1^-^, ckit^+^, CD34^int^, FcγRII/III^int^, Granulocyte Macrophage Progenitors (GMP) as Lin^-^, Sca1^-^, ckit^+^, CD34^int^, FcγRII/III^hi^. Hematopoietic stem cells (HSCs) were identified as Lin^-^, Sca1^+^, ckit^+^, CD48^-^ and CD150^+^.

**Supplemental Figure 5: Short-term propranolol treatment reduces cardiac injury and attenuates adverse remodeling following myocardial infarction (MI).**

**A**, Schematic figure depicting the study outline to test the effect of short term (ST) blockade of β1/β2 ARs on markers of cardiac injury and remodeling post-MI. **B**, Representative histological sections showing Ly6G (neutrophil) stain in left ventricle (LV) at 30 days post-MI and (**C**) its quantification (neutrophil counts/ field). **D**, Plasma cardiac troponin I levels measured at various time points post-MI. The inset shows quantification of AUC for each curve. **E**, Immunoblots showing the protein expression of α-smooth muscle actin (αSMA) and phospho (Thr202/Tr204)-ERK in LV homogenates along with total ERK and GAPDH (loading control). **F**, quantification of αSMA, normalized to GAPDH. **G**, quantification of phospho -ERK, normalized to total ERK and GAPDH. mRNA expression levels of collagen 1A1 (**H**), Collagen3A1 (**I**) and Fibronectin I (**K**) in LV homogenates (normalized to 18s). All data are means ± SEM. Statistical tests for **C** through **J**, 1-way ANOVA and Tukey’s post-hoc test.

**Supplemental Figure 6: Short-term inhibition of β2-ARs improves cardiac function and attenuates remodeling following MI.**

**A**, Schematic figure depicting the study outline to test the effect of short-term blockade of β1 ARs with metoprolol or β2 ARs with butaxamine post-MI. **B**, Vector diagrams showing the direction and magnitude of myocardial contraction. Three-dimensional regional wall displacement illustrations, demonstrating contraction (yellowish red) or relaxation (blue) of consecutive cardiac cycle results at 30 days post-MI. Measurement of left ventricular functional parameters by echocardiography including (**C**) Ejection fraction, (**D**) Cardiac output, (**E**) End Diastolic Volume and (**F**) End Systolic Volumes. Representative histological sections of left ventricle at 30 days post-MI showing (**G**) scar size by Masson’s trichrome staining and its (**H**) quantification represented as percentage whole heart (WH). Representative histological sections of left ventricle at 30 days post-MI showing picrosirius staining (**I**), a marker of cardiac fibrosis and (**J**) its quantification represented as percentage WH. All data are means ± SEM. n= 6-10 mice/ group. Statistical tests for **C** through **F**, Two-Way repeated measures ANOVA followed by a Bonferroni multiple comparison test. ^*^P <0.05 vs Vehicle and MI-MET groups. Statistical tests for **I** and **J,** 1-way ANOVA and Tukey’s post-hoc test (**^*^**P <0.05, **^**^**P <0.005. n.s, not significant).

**Supplemental Figure 7: Short-term inhibition of β1/ β2-ARs with propranolol suppresses neutrophil infiltration into the heart and improves cardiac function and remodeling following ischemia-reperfusion (IR) injury**.

**A**, Schematic figure depicting the study outline to test the effect of short-term (ST) blockade of β1/β2 ARs with propranolol in a model of IR injury. **B,** Representative flow cytometric plots and (**C**) quantification of neutrophils in the heart at 6 hours post reperfusion. **D,** Representative IHC images of left ventricle (LV) showing neutrophil abundance (Ly6G) and (**E**) their quantification at 6 hours post reperfusion. **F**, Plasma cardiac troponin I levels measured at various time points post-IR injury and (**G**) quantification of AUC for each curve. **H**, Vector diagrams showing the direction and magnitude of myocardial contraction. Three-dimensional regional wall displacement illustrations, demonstrating contraction (yellowish red) or relaxation (blue) of consecutive cardiac cycle results at 30 days post-IR. Measurement of left ventricular functional parameters by echocardiography including (**I**) Ejection fraction, (**J**) Cardiac output, (**K**) End Systolic Volume and (**L**) End Diastolic Volumes. Representative histological sections (Masson’s Trichrome stain, left panel) of LV at 30 days post-IR showing (**M**) scar size and (**N**) its quantification as percentage of whole heart (WH). Representative histological sections (Picrosirius stain, right panel) of LV at 30 days post-IR showing (**M**) cardiac fibrosis and (**O**) its quantification as percentage of WH. All data are means ± SEM. Statistical tests for **C, E, G** through **O**, 1-way ANOVA and Tukey’s post-hoc test (*P<0.05, **P<0.005, ***P<0.0005, ****P<0.0001, n.s, not significant). Statistical tests for **F** are Two-Way repeated measures ANOVA followed by a Bonferroni multiple comparison test (*P <0.05 vs Sham, ^#^P < 0.05 vs IR).
