## Supplementary figures and images for "β2 adrenergic receptors orchestrate neutrophil demargination and recruitment to the ischemic heart following myocardial infarction"

### Supplemental Figure -1

#### Slide 1
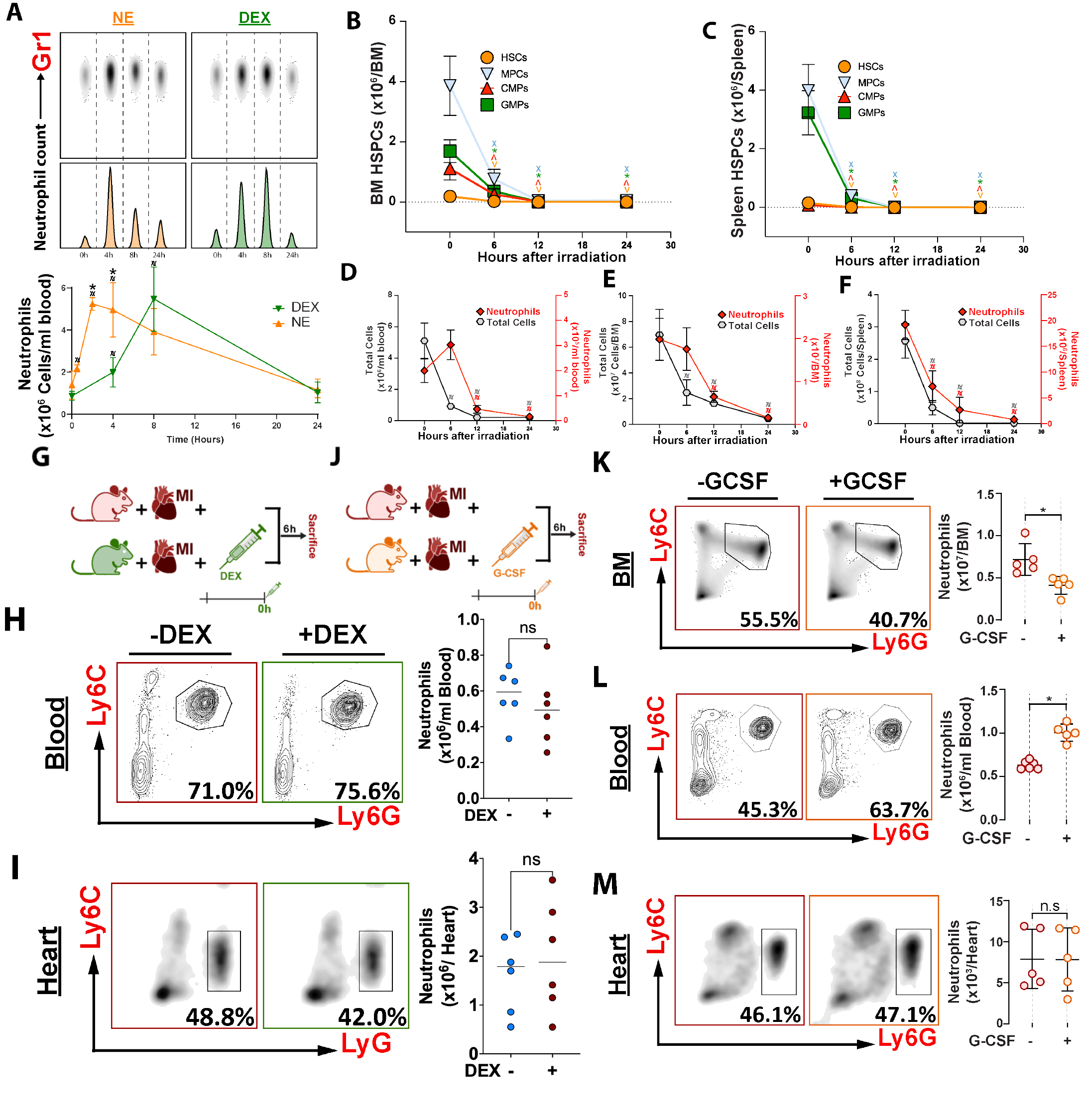

### Supplemental Figure -2

#### Slide 1
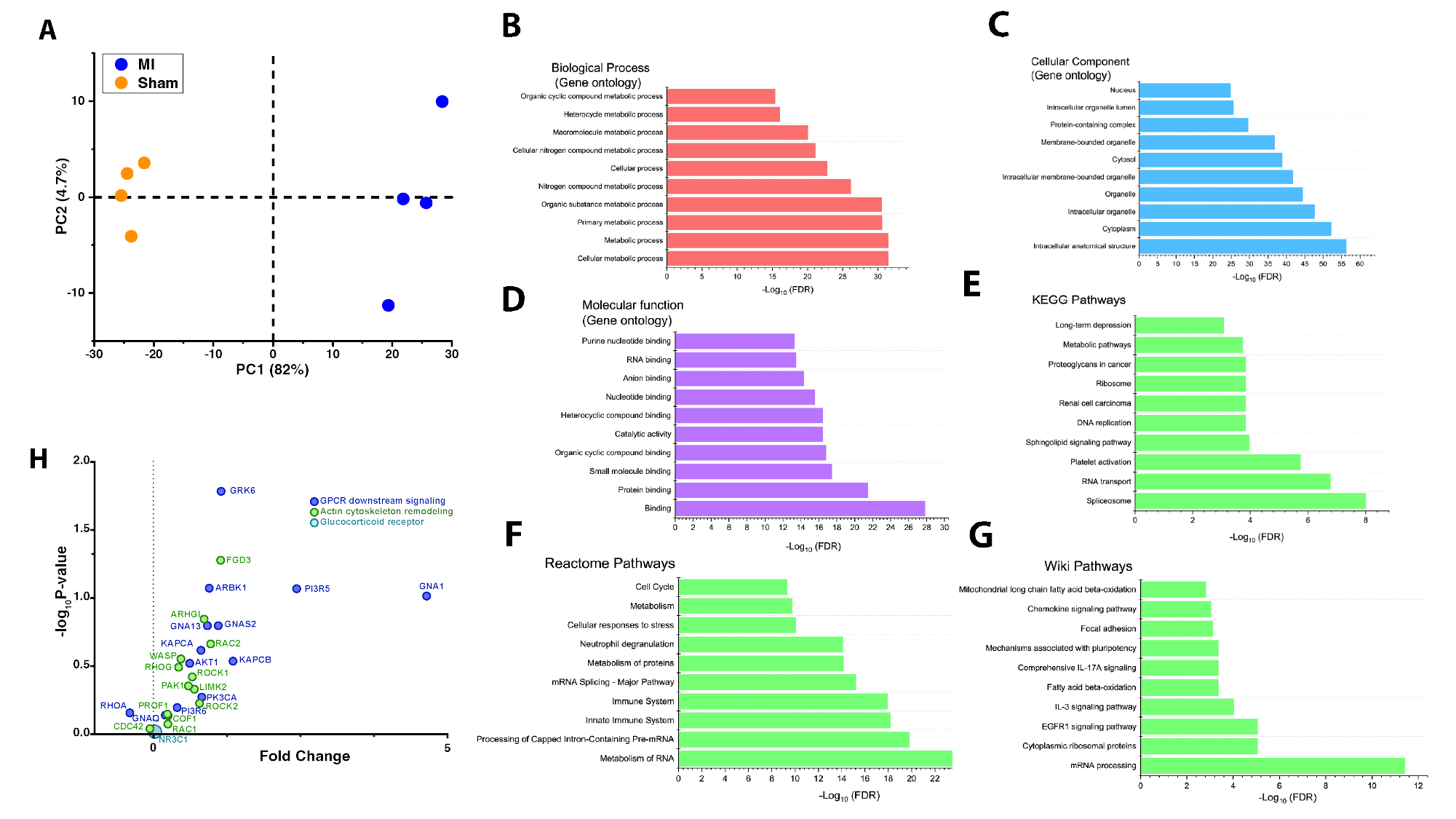

### Supplemental Figure -3

#### Slide 1
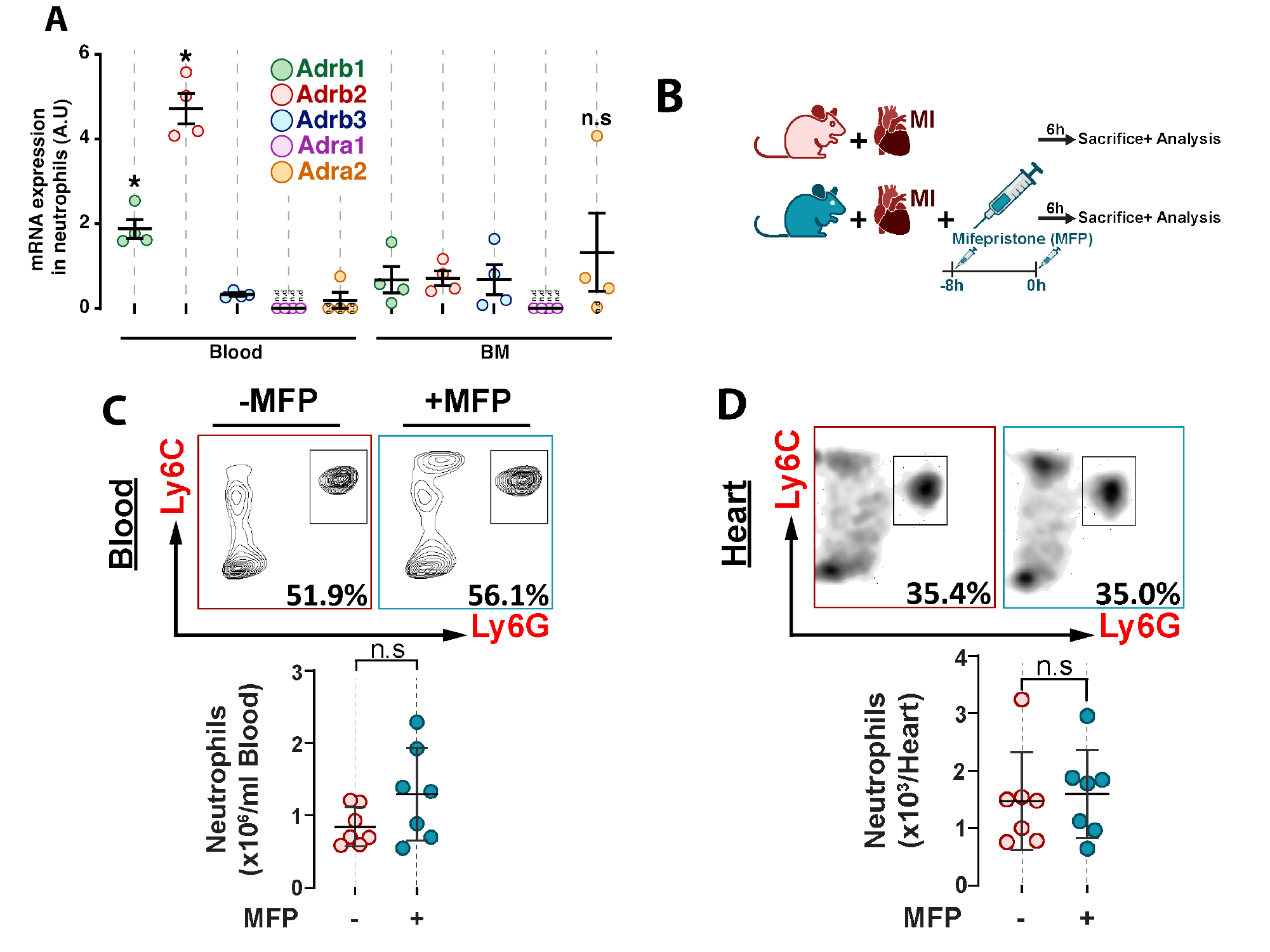

### Supplemental Figure -4

#### Slide 1
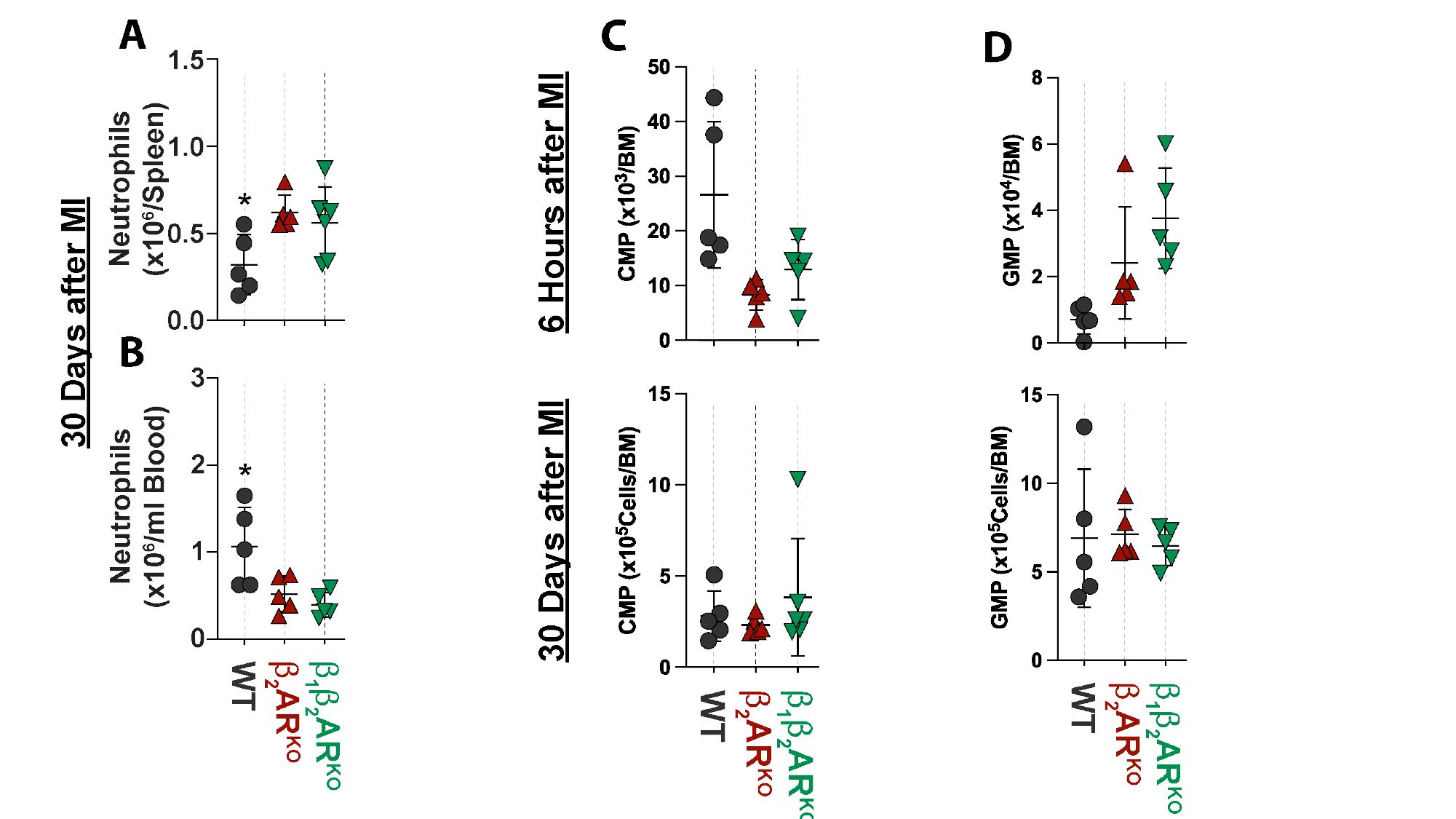

### Supplemental Figure -5

#### Slide 1
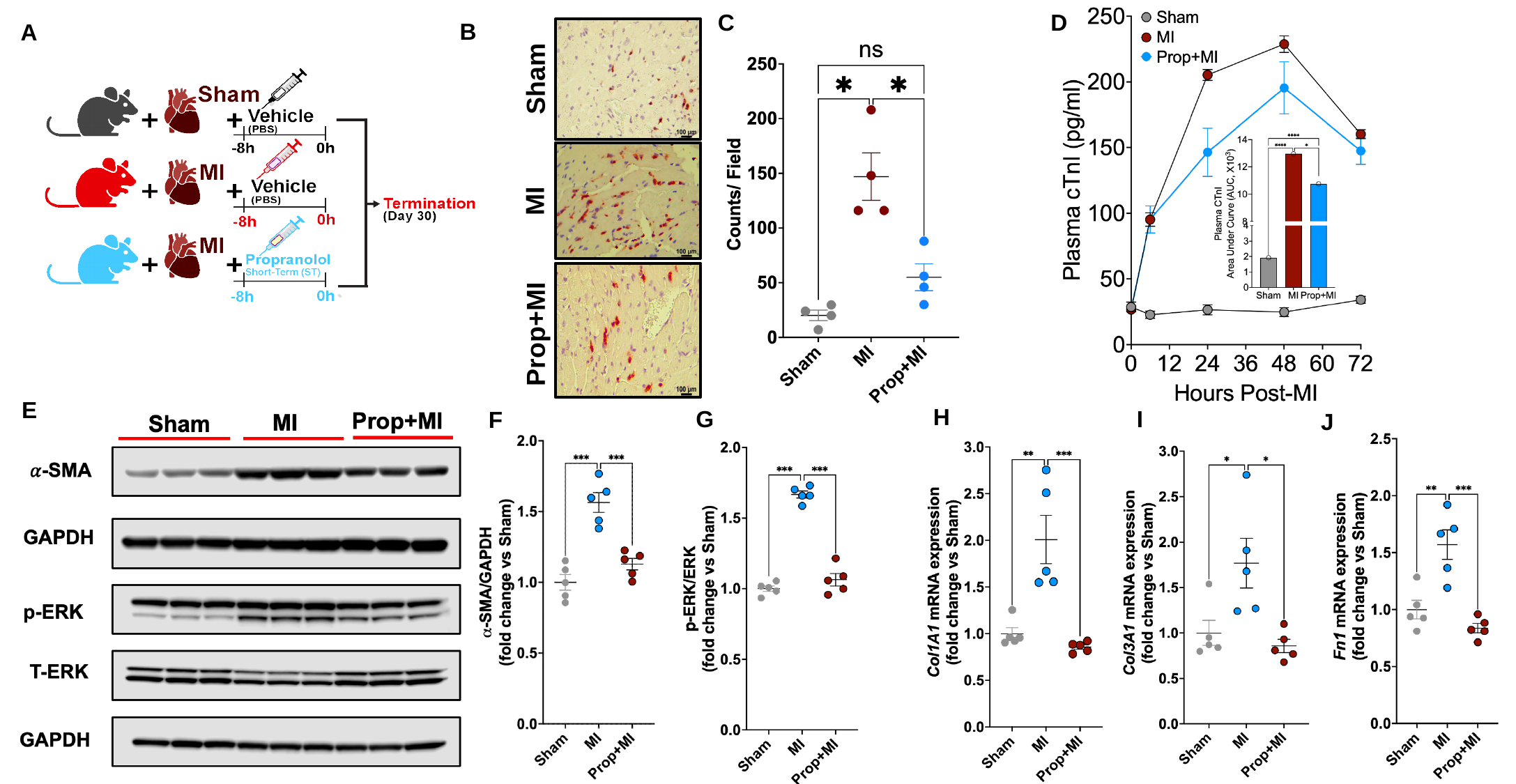

C
D
B
A
Sham
MI
Prop+MI
E
H
G
I
F
J

### Supplemental Figure -6

#### Slide 1
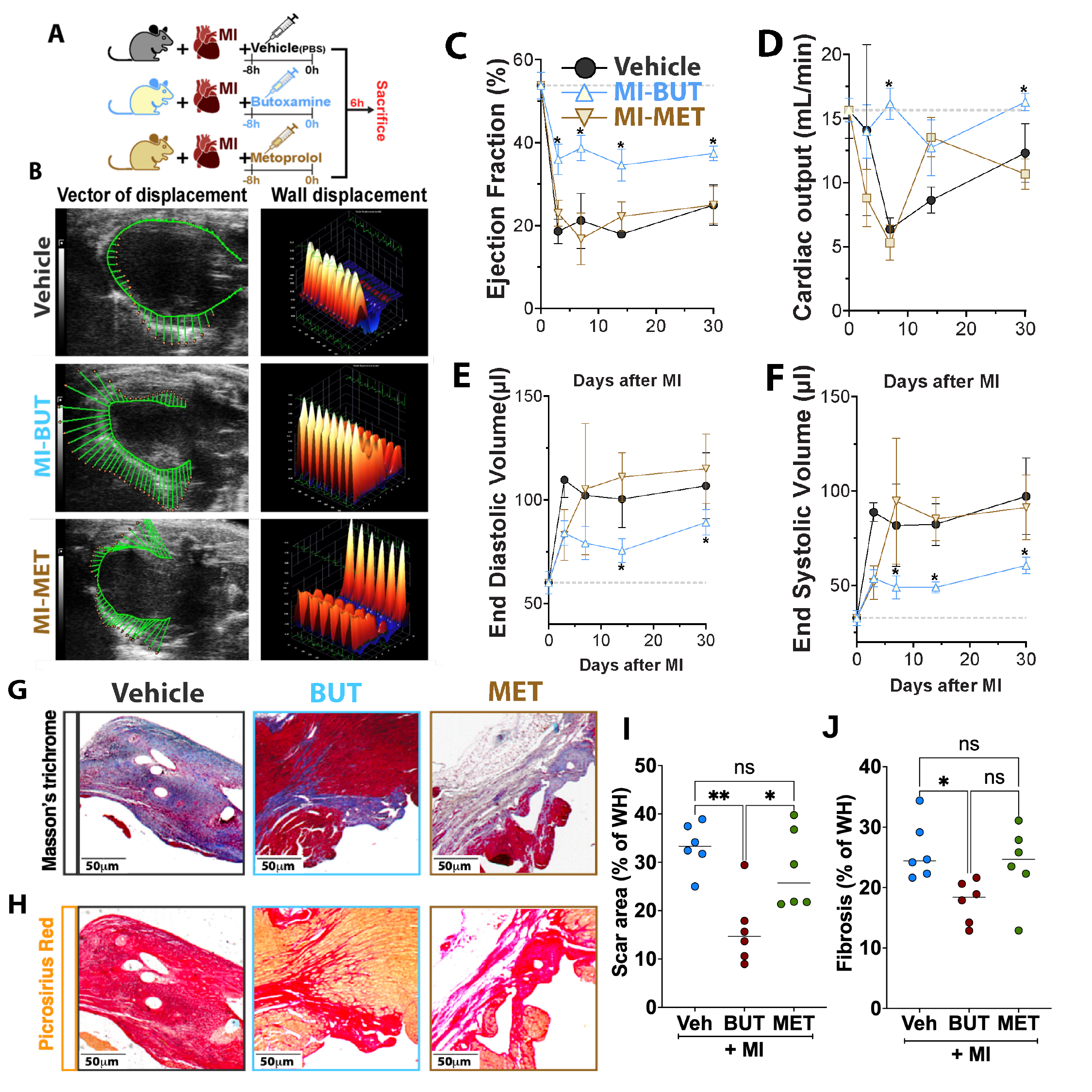

J
I

### Supplemental Figure -7

#### Slide 1
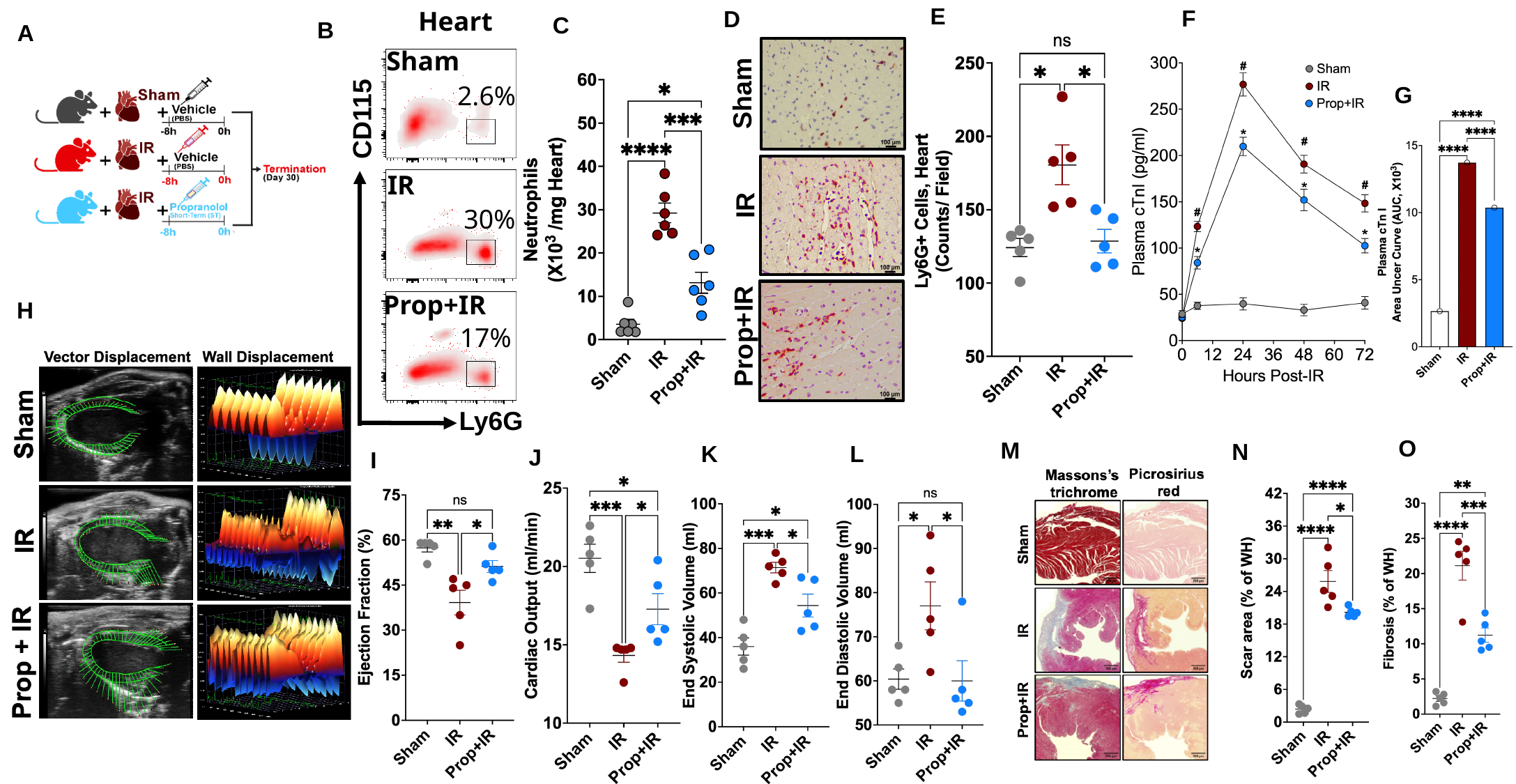

Heart
E
F
D
C
B
A
Sham
CD115
Ly6G
2.6%
30%
17%
IR
Prop+IR
Sham
IR
Prop+IR
G
H
O
M
N
K
L
J
I
